# Unveiling the signaling network of FLT3-ITD AML improves drug sensitivity prediction

**DOI:** 10.1101/2023.06.22.546072

**Authors:** Sara Latini, Veronica Venafra, Giorgia Massacci, Valeria Bica, Simone Graziosi, Giusj Monia Pugliese, Marta Iannuccelli, Filippo Frioni, Gessica Minnella, John Donald Marra, Patrizia Chiusolo, Gerardo Pepe, Manuela Helmer-Citterich, Dimitrios Mougiakakos, Martin Boettcher, Thomas Fischer, Livia Perfetto, Francesca Sacco

**Author notes:** These authors have contributed equally to this work (Sara Latini, Veronica Venafra).

## Abstract

Currently, the identification of patient-specific therapies in cancer is mainly informed by personalized genomic analysis. In the setting of acute myeloid leukemia (AML), patient-drug treatment matching fails in a subset of patients harboring atypical internal tandem duplications (ITDs) in the tyrosine kinase domain of the FLT3 gene. To address this unmet medical need, here we develop a systems-based strategy that integrates multiparametric analysis of crucial signaling pathways, and patient-specific genomic and transcriptomic data with a prior-knowledge signaling network using a Boolean-based formalism. By this approach, we derive personalized predictive models describing the signaling landscape of AML FLT3-ITD positive cell lines and patients. These models enable us to derive mechanistic insight into drug resistance mechanisms and suggest novel opportunities for combinatorial treatments. Interestingly, our analysis reveals that the JNK kinase pathway plays a crucial role in the tyrosine kinase inhibitor response of FLT3-ITD cells through cell cycle regulation. Finally, our work shows that patient-specific logic models have the potential to inform precision medicine approaches.

## Introduction

In the era of precision medicine, comprehensive profiling of malignant tumor samples is becoming increasingly time- and cost-effective in clinical ecosystems (De Maria Marchiano et al., 2021; Tsimberidou et al., 2020). While a growing number of genotype-tailored treatments have been approved for use in clinical practice (Krzyszczyk et al., 2018; Scheetz et al., 2019), the success of targeted therapies is limited by frequent development of drug resistance mechanisms that lead to therapy failure and portend a dismal patient prognosis (Mansoori et al., 2017; Sabnis and Bivona, 2019; Vander Velde et al., 2020; Vasan et al., 2019). Drug combinations are currently under investigation as a potential means of avoiding drug resistance and achieving more effective and durable treatment responses.

As the number of possible combinations increases exponentially with the number of drugs available, it is impractical to test for potential synergistic properties among all available drugs using empirical experiments alone. Computational approaches that can predict drug synergy, including Boolean logic models, are crucial in guiding experimental approaches for discovering rational drug combinations. In the Boolean model, a biological process or pathway of interest is modeled in the form of a signed and direct graphic with edges representing the regulatory relationship (activating or inhibitory) between the nodes (proteins). Logical operators (AND, OR, and NOT), are then employed to dynamically describe how the signal is integrated and propagated in the system over time to reach a terminal state. These states can be associated with cellular processes such as apoptosis and proliferation (Calzone et al., 2022). Once optimized, these models offer the ability to test for the effect of perturbation of the nodes on the resulting phenotype (e.g., *in silico* knockout), allowing us to generate novel hypotheses and to predict the efficacy of novel drug combinations (Hemedan et al., 2022; Le Novère, 2015; Montagud et al., 2022; Schwab et al., 2020; Wang et al., 2012).

Among the different computational methods available, in the present study, we utilized CellNOptR (Terfve et al., 2012) to implement an integrated strategy that combines prior-knowledge signaling networks (PKN) with multiparametric analysis and Boolean logic modeling. We applied this approach to generate genotype-specific predictive models of AML patients with differing sensitivities to drug treatments. Specifically, we focused on a subset of AML patients with internal tandem duplication (ITDs) in the FLT3 receptor tyrosine kinase. FLT3-ITD, one of the most common driver mutations in AML, occurs in exons 15 and 16, which encode the juxtamembrane domain (JMD) and the first tyrosine-kinase (TKD1) domain, and results in constitutive activation. We and others have demonstrated that the location (insertion site) of the ITD is a crucial prognostic factor: treatment with the recently FDA-approved multi-kinase inhibitor Midostaurin and standard frontline chemotherapy has a significant beneficial effect only in patients carrying the ITDs in the JMD domain, whereas no beneficial effect has been shown in patients carrying ITDs in the TKD region (Rücker et al., 2022; Pugliese et al., 2023; Massacci et al., 2023). Moreover, our group and others have demonstrated the differences underlying Tyrosine Kinase Inhibitor (TKI) sensitivity are related to a genotype-specific rewiring of the involved signaling networks.

In the present study, we applied a newly developed integrated approach to construct predictive logic models of cells expressing FLT3^ITD-TKD^ and FLT3^ITD-JMD^. These models revealed that JNK plays a crucial role in the TKI response of FLT3-ITD cells through a cell cycle-dependent mechanism, in line with our previous findings (Massacci et al., 2023; Pugliese et al., 2023). Additionally, we integrated patient-specific genomic and transcriptomic data with cell line-derived logic models to obtain predictive personalized mathematical models with the aim of proposing novel patient-individualized anti-cancer treatments.

## Results

### The experimental strategy

In the treatment of cancer, molecular-targeted therapies often have limited effectiveness, as tumors can develop resistance over time. One potential solution to this problem is the use of combination therapy, for which data-driven approaches are valuable in identifying optimal drug combinations for individual patients. To identify novel genotype-specific combinatorial anti-cancer treatments in AML patients with FLT3-ITD, we employed a multidisciplinary strategy combining multiparametric analysis with literature-derived causal networks and Boolean logic modeling. Our experimental model consisted of hematopoietic Ba/F3 cells stably expressing the FLT3 gene with ITDs insertions in the JMD domain (aa 598) or in the TKD1 (aa 613) region (henceforth “FLT3^ITD-JMD^” and “FLT3^ITD-TKD^” cells, respectively). As previously demonstrated, cells expressing FLT3^ITD-TKD^ (“resistant model”) have significantly decreased sensitivity to TKIs, including the recently registered FLT3-TKI Midostaurin, compared with FLT3^ITD-JMD^ cells (“sensitive model”) (Massacci et al., 2023; Pugliese et al., 2023). Our approach (schematized in Figure 1) is summarized as follows:

STEP 1: The first step in our strategy aimed at providing a detailed description of FLT3-ITD-triggered resistance mechanisms. To this end, we carried out a curation effort and mined our in-house resource, SIGNOR (Lo Surdo et al., 2023), to build a prior-knowledge network (PKN) recapitulating known signaling pathways downstream of the FLT3 receptor. The PKN integrates information obtained in different cellular systems under distinct experimental conditions (**Fig. 1, panel A**).

STEP 2: Using the PKN, we selected 14 crucial proteins, which we refer to as ‘sentinel proteins’, whose protein activity was emblematic of the cell state downstream of FLT3. Thus, by performing a multi-parametric analysis, we measured the activity status of the sentinel proteins under 16 different perturbation conditions in TKIs sensitive and resistant cells to generate the training dataset (**Fig. 1, panel B**).

STEP 3: We employed the CellNOptR tool to optimize the PKN using the training data. Two genotype-specific predictive models were generated that best reproduced the training dataset (**Fig. 1, panel C**).

STEP 4: Using the optimized model, we performed an *in silico* knock-out screen involving the suppression of multiple crucial nodes. Novel combinatorial treatments were predicted according to the induction of apoptosis in TKI-sensitive and TKI-resistant cells (**Fig. 1, panel D**).

STEP 5: The predictive performance of the two models was validated *in vitro*, and *in silico* in two independent publicly available datasets. The clinical impact of our models was assessed in a cohort of 14 FLT3-ITD positive AML patients (**Fig. 1, panel E**).

**Figure 1.**
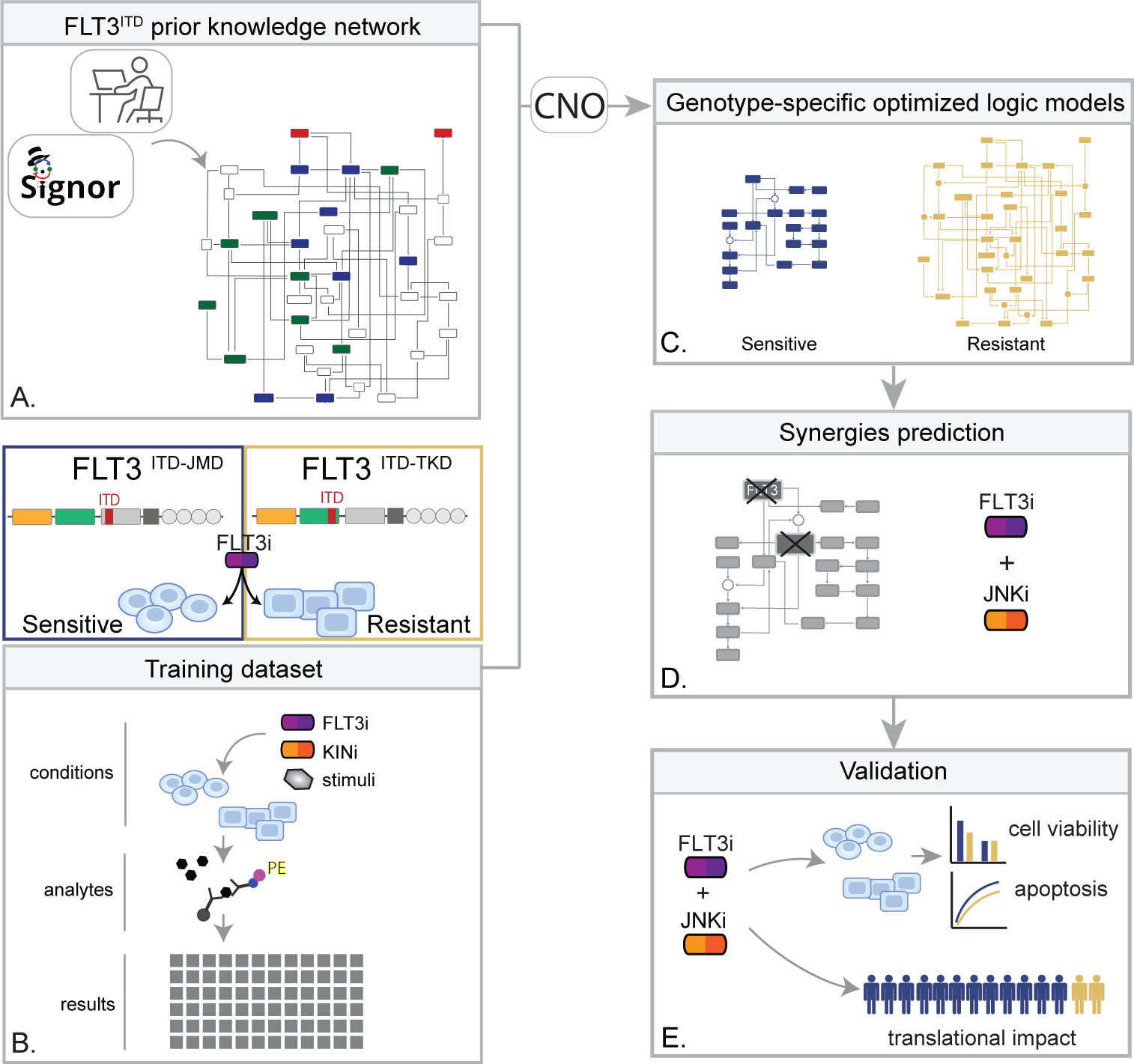
Summary of the experimental strategy. **A**) Manual curation of FLT3-ITD-specific Prior Knowledge Network (PKN). **B**) Multiparametric analysis of signaling perturbations in TKI sensitive and TKI resistant cells. **C**) Model generation through the CellNOptR tool. **D**) Prediction of combinatorial treatments restoring TKI sensitivity. **E**) *In vitro* validation of novel combinatorial treatments. **F**) *In silico* prediction of co-treatment outcome in AML patients.

### Generation of FLT3-ITD prior-knowledge signaling network

The first step in the application of our pipeline consisted of the creation of the Prior Knowledge Network (PKN), a static and genotype-agnostic map recapitulating the signaling pathways deregulated over AML tumor development and progression (**Fig. S1**). To create the PKN, we embarked on a curation effort aimed at describing the molecular mechanisms or causal relationships connecting three crucial receptors responsible for sustaining the proliferative and survival pathways in AML (FLT3, TNFR, and IGF1R), to downstream events (i.e., apoptosis and proliferation). Gathered data were captured using our in-house developed resource, SIGNOR, and made freely accessible to the community for reuse and interoperability, in compliance with the FAIR principles (Wilkinson et al., 2016). Briefly, SIGNOR (https://signor.uniroma2.it) is a public repository that captures more than 35K causal interactions (up/down regulations) among biological entities and represents them in the form of a direct and signed network (Lo Surdo et al., 2023). This representation format makes it particularly suitable for the implementation of Boolean logic modeling approaches. The so-obtained pre-PKN included 76 nodes and 193 edges, the nodes representing proteins, small molecules, stimuli, and phenotypes, and the edges depicting the directed interactions between the nodes (**Table S1**).

As little is known about the specific signaling pathways downstream of the non-canonical FLT3^ITD-TKD^, we enriched the pre-PKN, deriving new edges from cell-specific experimental data of both FLT3^ITD-TKD^ and FLT3^ITD-JMD^ expressing cell lines (see methods, **Fig. 2A**). This refined PKN recaps the FLT-ITD downstream signaling and served as the basis for the model optimization.

**Figure 2.**
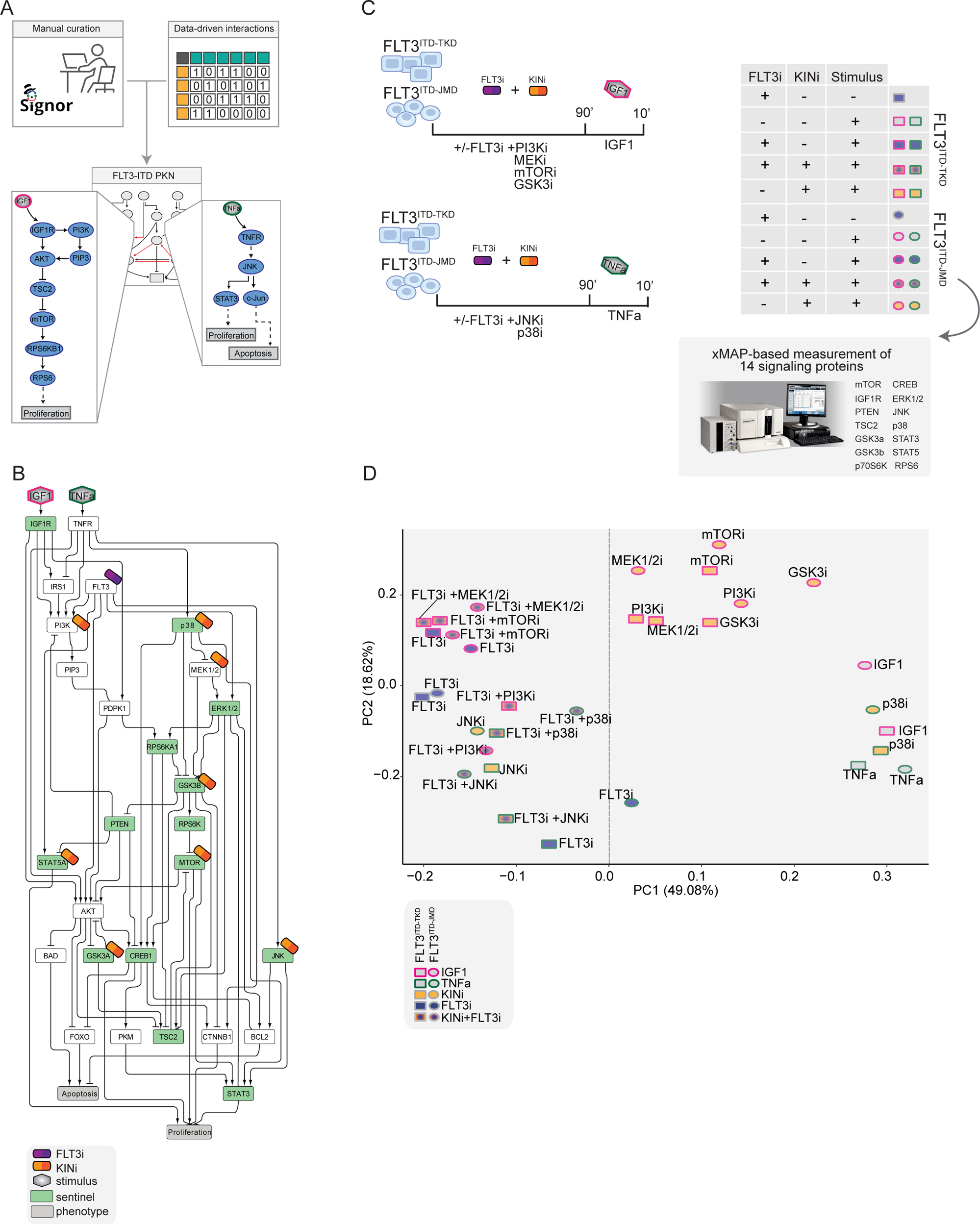
Generation of the training dataset. **A)** Schematic representation of the FLT3-ITD PKN manual curation, integration of data-driven edges, and manual integration of RTKs pathways involved in AML. **B)** Schematic representation of the experimental design: FLT3^ITD-JMD^ and FLT3^ITD-TKD^ cells were cultured in starvation medium (w/o FBS) overnight and treated with PI3Ki, MEKi, mTORi, and GSK3i, JNKi and p38i, in presence or absence of the FLT3i Midostaurin for 90 minutes. Then, the cells were stimulated either with IGF1 or TNFα for 10 minutes. Control cells were starved and treated with Midostaurin for 90 minutes. After treatment, samples were collected, and cell lysates were analyzed with an xMAP-based assay through the MagPix instrument. Per each experimental condition, we measured the phosphorylation levels of 14 sentinels. **C)** Network representation of a compressed PKN that shows the essential pathways monitored through the perturbation experiment. The perturbed nodes are tagged with a drug icon, and the measured nodes are colored green. **D)** Principal Component Analysis (PCA) of FLT3^ITD-JMD^ and FLT3^ITD-TKD^ cells in the multiparametric analysis. Each point represents a different experimental condition.

### Multiparametric analysis of TKI-resistant and sensitive FLT3-ITD cells

To clarify the cooperative and antagonistic interactions among FLT3 inhibition and complementary therapeutic strategies, we generated a cue–sentinel–response multiparametric dataset (**Table S2-4**). Generally, MAPKs, PI3K-AKT-mTOR, and STATs are the main pathways downstream of FLT3, IGF1R, and TNFR; we selected 6 key kinases in the FLT3-ITD PKN, and we perturbed their activity with small molecule inhibitors in presence or in absence of the FLT3i Midostaurin. We added the cytokines as stimuli to fully activate the RTKs included in our network (**Fig. 2B**). Specifically, we treated sensitive and resistant cells with either PI3Ki, mTORi, MEKi, or GSKi +/− Midostaurin for 90 minutes and we added IGF1 for the last 10 minutes. Parallely, we treated the cells with p38i or JNKi +/− Midostaurin for 90 minutes and added TNFα for the last 10 minutes (**Fig. 2C**). Overall, we subjected our cell lines to 16 experimental conditions (listed in Methods, **Table S2**) and in each of them we measured the signaling perturbations. As sentinels of the signaling activity response, we measured in triplicate the activity states of 14 crucial proteins (**Fig. 2B-C**) based on their phosphorylation status (mTOR, CREB1, IGF1R, PTEN, GSK3a, GSK3b, STAT3, STAT5, TSC2, p70S6K, RPS6, JNK, p38, ERK1/2).

Briefly, the biological replicates displayed Pearson correlation coefficients ranging between 0.75-1 (**Fig. S2A-B**). Overall, the observed modulation of the readouts was consistent with the experimental evidence reported in the literature (**Fig. S2C,** black squares in the heatmap). For each sentinel protein, we employed combinations of inhibitors and stimuli to probe the full spectrum of protein activity, ranging from the minimum (inhibitor treatment) to the maximum (stimulus exposure). The data were normalized in the 0 to 1 range using a Hill function. In this way, the fully active sentinel value was = 1, and the inhibited value = 0 (**Fig. S3**).

Principal component analysis (PCA) (**Fig. 2D**) and unsupervised hierarchical clustering (**Fig. S2D**) showed that the activity level of sentinel proteins stratified cells according to both FLT3 activation status (component 1: presence vs absence of FLT3i) and cytokine stimulation (component 2: IGF1 vs TNFα). Of note, among all the KINi-treated conditions, only the JNKi treatment groups with the FLT3i treated samples in both cell lines. On the contrary, the activation profile of these 14 sentinel proteins was not able to distinguish cells according to the distinct FLT3-ITD insertion sites (circles=FLT3^ITD-JMD^ and squares=FLT3^ITD-TKD^) (**Fig. 2D**). Interestingly, the unsupervised hierarchical clustering of the 14 analytes revealed different groups according to the pathway proximity of the nodes (e.g., JNK-p38; p70S6K-RPS6; STAT3-STAT5) or according to their regulatory role (e.g., GSK3a/b, PTEN and TSC2, acting as negative regulators, cluster together) (**Fig. S2E**). Together, these observations suggest that the characterization of the genotype-dependent rewiring of signaling pathways cannot be obtained by simply looking at single proteins in our multiparametric dataset, but rather requires a modeling approach.

### Generation of FLT-ITD optimized logic models

CellNOptR was used to derive biologically relevant information from our dataset and generate FLT3-ITD-specific predictive models. Boolean logic models were optimized by maximizing the concordance between the PKN and our cue–sentinel–response multiparametric training dataset (**Fig. 3A**).

**Figure 3.**
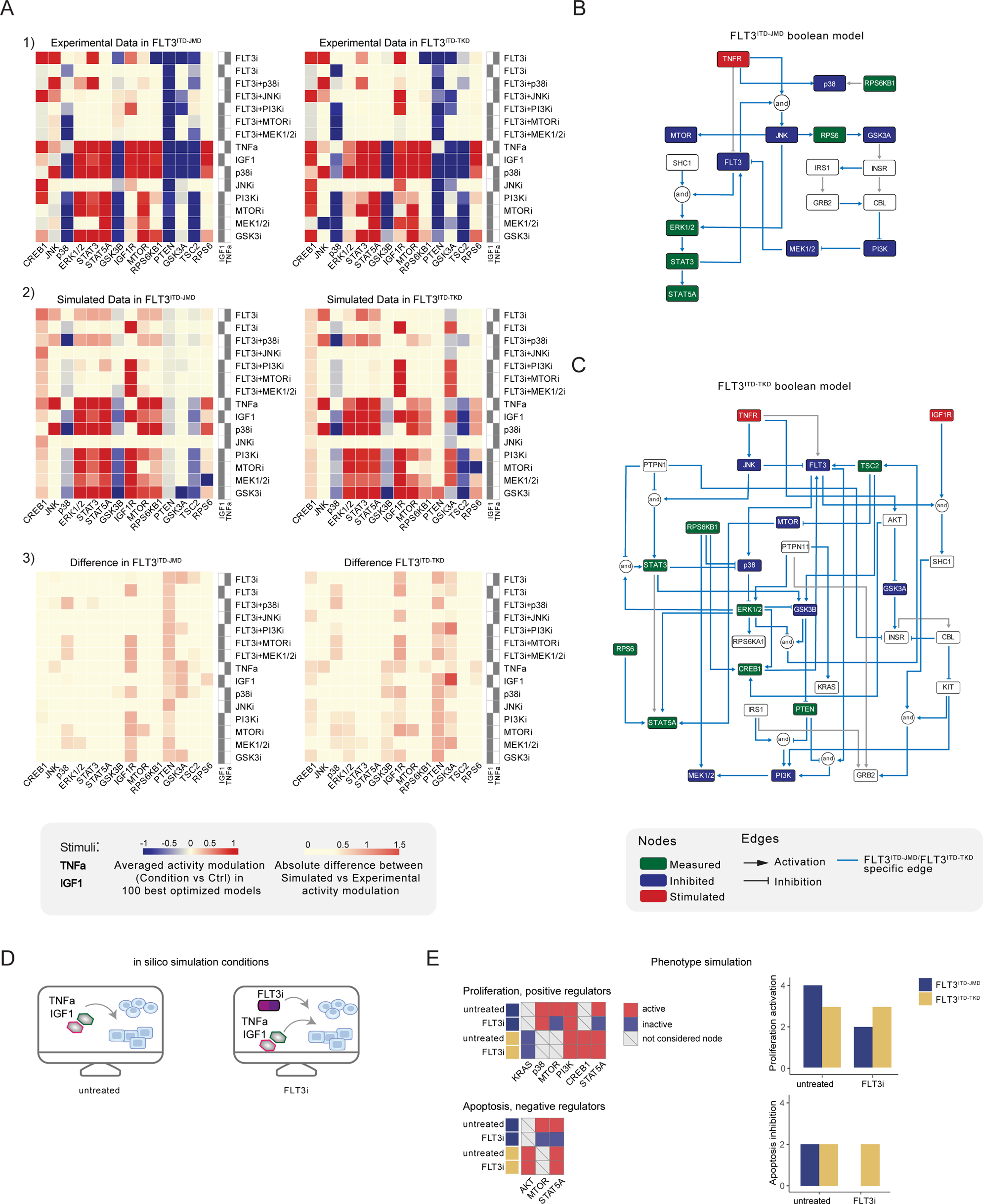
Optimized Boolean models recapitulate the different sensitivity of FLT3^ITD-JMD^ and FLT3^ITD-TKD^ cells to TKI. **A)** Color-coded representations of the experimental activity modulation (T90 – T0) of sentinel proteins used to train the two Boolean models (*upper panel*) and the average prediction of protein activities in the family of 100 best models (*central panel*). The protein activity modulation ranges from −1 to 1 and is represented with a gradient from blue (inhibited) to red (activated). The absolute value of the difference between experimental and simulated protein activity modulation (*lower panel*) is reported as a gradient from light yellow (error < 0.5) to red (1.85). **B-C)** FLT3^ITD-JMD^ (**B**) and FLT3^ITD-TKD^ (**C**) high-confidence Boolean models. Perturbed proteins in the experimental setup are marked in red or green if inhibited or stimulated, respectively. Sentinel proteins are reported in blue. The edges’ weight represents their frequency in the family of 100 models and only the high-confidence ones (frequency > 0.4) are reported. Orange edges are cell-specific links. **D)** Cartoon of the *in silico* conditions simulated to analyze the different TKI sensitivity of the FLT3^ITD-JMD^ and FLT3^ITD-TKD^ Boolean models. Untreated condition: TNFα+IGF1; FLT3i: TNFα+IGF1+FLT3 inhibition. **E)** Heatmaps (left) report the activation level of positive and negative phenotype regulators present in the two Boolean models. Bar plots (right) showing the proliferation activation and apoptosis inhibition levels in untreated and FLT3i conditions in the steady states of FLT3^ITD-JMD^ (blue) and FLT3^ITD-TKD^ (yellow) Boolean models.

In the first step, CellNOptR preprocesses the PKN (**Fig. S1**) and translates it into logical functions (scaffold model). As previously described (Sacco et al., 2012), the preprocessing consists of three phases: i) *compression*, in which unmeasured and untargeted proteins, as well as linear cascades of undesignated nodes, are removed; ii) *expansion*, in which the remaining nodes are connected to the upstream regulators with every possible combination of OR/AND Boolean operators; and iii) *imputation,* in which the software integrates the scaffold model with regulations function inferred without bias from the training dataset.

Using this strategy, we obtained two FLT3-ITD specific PKNs, accounting for 206 and 208 nodes and 756 and 782 edges, for FLT3^ITD-JMD^ and FLT3^ITD-TKD^ respectively. The variation in the node count between these two PKNs results from the inclusion of a different number of AND Boolean operators during the *expansion* step, while the difference in edge numbers is primarily due to different variations in the data, leading to distinct edge connections in the *imputation* step.

In the second step, causal paths and Boolean operators from the scaffold models were filtered to best fit the experimental context (see Methods). Briefly, for each cell line, we trained the software with our normalized cue–sentinel–response multiparametric dataset to generate a family of 1000 optimized Boolean models, and we retained the top 100 performing models (**Fig. S4A**).

To qualitatively assess the robustness and reliability of the selected models, we compared the average activity modulation of the individual sentinel proteins with experimentally observed readouts (**Fig. 3A, panel 1-2**).

Since the performance of the model strongly depends on the topology of the PKN, we performed several rounds of PKN check and adjustment, and, in each round, the entire process was iterated until the simulation provided the best fit of the available data (**Fig. S4B-C**). As shown in Fig. 3A, panel 3, the fit between simulated and experimental data was generally higher in the FLT3^ITD-JMD^ model, which has been more extensively characterized by the scientific community than the FLT3^ITD-TKD^ system. For each cell line, we selected the model with the lowest error (see Methods) between experimental and simulated data in the two cell lines (best model) (**Fig. 3B-C**). Interestingly, the two Boolean models display a different structure, and most of the interactions are cell-specific (blue edges), with only a few edges shared among the two networks (e.g., TNFR-FLT3, STAT3-STAT5A, p70S6K-p38, etc.). The architectural differences between the models demonstrate a profound rewiring of the signal downstream of FLT3 as a result of the different locations of the ITD.

### Evaluating the predictive power of FLT3-ITD logic models

Thus, we first took advantage of the publicly available quantitative phosphoproteomics dataset to independently validate our models. To this aim, we computed the steady state of the two models in “untreated” and “FLT3i” conditions (**Fig. S4D**). Briefly, the *untreated* condition represents the tumor state, here the pro-survival receptors (FLT3, IGF1R, and TNFR) are set constitutively active and assigned a Boolean value of 1. In the *FLT3i* condition (Midostaurin administration), the FLT3 receptor is inhibited and associated with a Boolean value of 0, whereas IGF1R and TNFR remain constitutively active to reflect the environmental background that sustains tumor growth and proliferation (**Fig. 3D**). Given these two initial conditions (*untreated* vs *FLT3i*), we carried out a synchronous simulation (Schwab et al., 2020) to compute the evolution of the two models. Next, we compared the steady state of our model upon FLT3 inhibition with the phosphoproteomic data describing the modulation of 16,319 phosphosites in FLT3-ITD Ba/F3 cells (FLT3^ITD-TKD^ and FLT3^ITD-JMD^) upon quizartinib (AC220) treatment (Massacci et al., 2023). The activation status of the nodes in the two generated models is highly comparable with the level of regulatory phosphorylations reported in the reference dataset, supporting our models (**Fig. S5A**). Next, we aimed to assess whether the newly generated FLT3^ITD-JMD^ and FLT3^ITD-TKD^ Boolean models could recapitulate *in silico* the modulation of apoptosis and proliferation upon inhibition of FLT3 and other druggable nodes of our models. First, to functionally interpret the results of the simulations, for each network, we extracted key regulators of ‘apoptosis’ and ‘proliferation’ hallmarks from SIGNOR. To this aim, we applied our recently developed ProxPath algorithm, a graph-based method able to retrieve significant paths linking the nodes of our two optimized models to proliferation and apoptosis phenotypes ((Iannuccelli et al., 2023), see Methods) (**Table S1, Fig. 3E, left panels, Fig. S4D**). Then, integrating the signal of their key regulators (see Methods), we were able to derive the ‘proliferation’ and ‘apoptosis inhibition’ levels upon each initial condition.

Importantly, our strategy demonstrated that FLT3^ITD-JMD^ and FLT3^ITD-TKD^ Boolean models were able to recapitulate the different TKI sensitivity of FLT3-ITD cells (**Fig. 3E, right panels**) (Massacci et al., 2023; Pugliese et al., 2023).

Moreover, by taking advantage of the Beat AML program, which provides *ex vivo* drug sensitivity screening data of 134 FLT3^ITD-JMD^ AML patients, we validated the prediction power of our models by comparing our *in silico* results with the in vitro IC50 values measured upon RTKs inhibition (**Fig. S5B-C**). We observed some discrepancies between model’s prediction and patients’ data for PI3K inhibition (probably due to missing connections in our cell-specific model) while FLT3, mTOR, JNK and p38 treatments outcomes in patients were successfully predicted by our models.

### Identification of novel combinatorial treatments reverting TKI resistance

As per their intrinsic nature, the two optimized Boolean logic models have predictive power and can be used to simulate *in silico* novel combinatorial treatments reverting drug resistance of FLT3^ITD-TKD^ cells (**Fig. 4A**).

**Figure 4.**
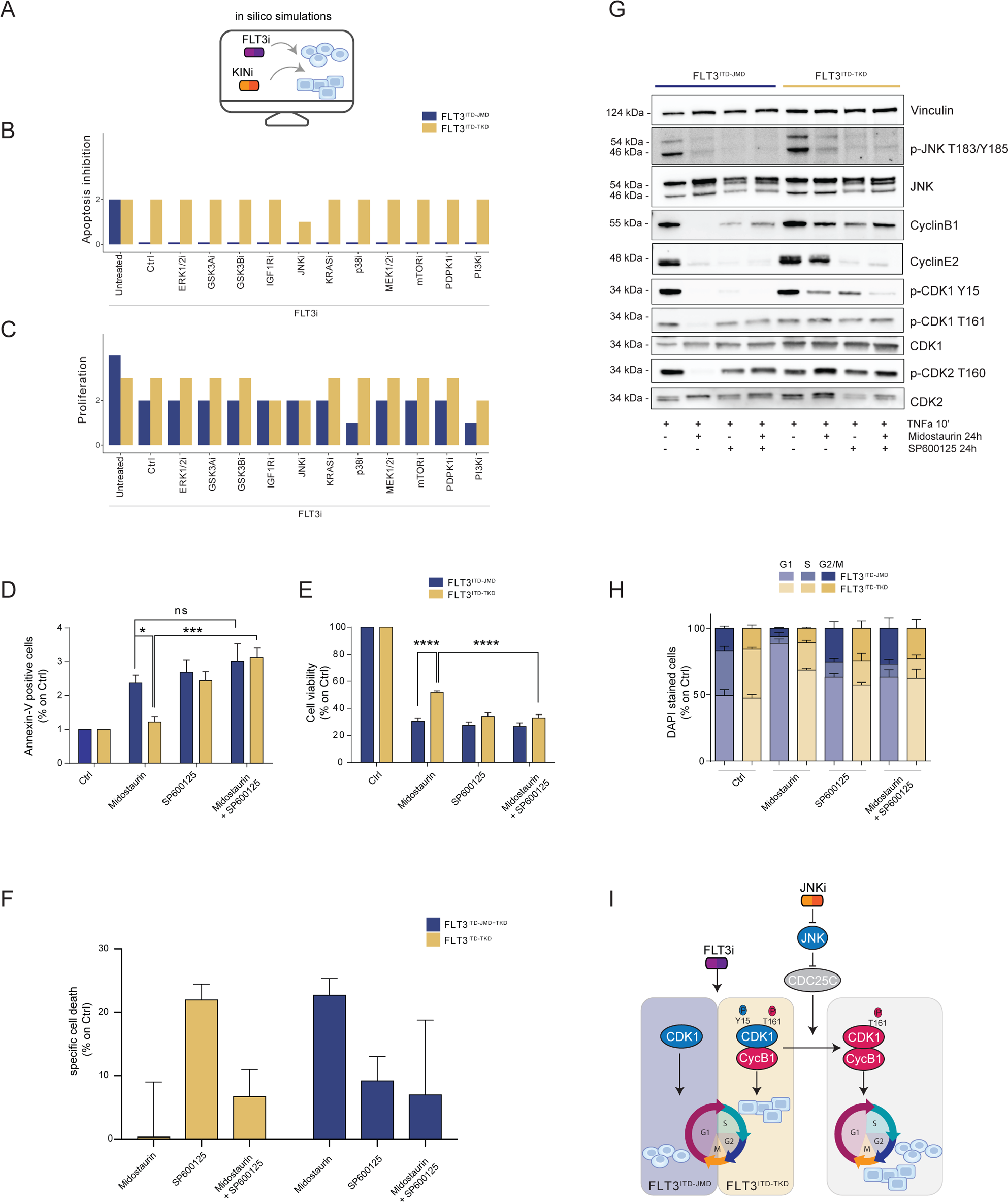
In silico simulation of the FLT3^ITD-TKD^ logic model allows the prediction of novel combinatorial treatment reverting TKI resistance. **A)** Cartoon of the *in silico* simulation conditions. **B**-**C)** Bar plot showing the *in silico* simulation of proliferation activation (**B**) and apoptosis inhibition (**C**) levels in untreated and FLT3i conditions in combination with knock-out of each of 10 crucial kinases in FLT3^ITD-JMD^ (blue) and ^-TKD^ (yellow) cells. **D-E**) In FLT3^ITD-JMD^ (blue) and -TKD (yellow) cells treated with 100nM Midostaurin and/or 10uM SP600125 (JNK inhibitor) for 24h, the percentage of Annexin V positive cells (**D**) and the absorbance values at 595nm (**E**), normalized on control condition, are reported in bar plots. **F)** Primary samples from AML patients with the FLT3^ITD-TKD^ mutation (n=2, yellow bars) or the FLT3^ITD-JMD/TKD^ mutation (n=3, blue bars) were exposed to Midostaurin (100nM, PKC412), and JNK inhibitor (10µM, SP600125) for 48 hours, or combinations thereof. The specific cell death of gated AML blasts was calculated to account for treatment-unrelated spontaneous cell death. The bars on the graph represent the mean values with standard errors. **G)** In FLT3^ITD-JMD^ (blue) and FLT3^ITD-TKD^ (yellow) cells treated with 100nM Midostaurin and/or 10uM SP600125, followed by 10’ of TNFα 10ng/ml, the protein levels of phospho-JNK (T183/Y185), JNK, phospho-CDK1 (Y15), phospho-CDK1 (T161), CDK1, phospho-CDK2 (T160), CDK2, CyclinB1, CycinE2, and Vinculin were evaluated by western blot analysis. **H)** Cytofluorimetric cell cycle analysis of DAPI stained FLT3^ITD-JMD^ (blue) and FLT3^ITD-TKD^ (yellow) cells treated with 100nM Midostaurin and/or 10uM SP600125 (JNK inhibitor) for 24h. **I)** Cartoon of the molecular mechanism proposed for FLT3^ITD-JMD^ and FLT3^ITD-TKD^ cells.

Thus, we performed a targeted *in silico* approach in FLT3^ITD-TKD^ and FLT3^ITD-JMD^ cells, by simulating the levels of apoptosis and proliferation, upon combinatorial knockout of FLT3 and one of the following key druggable kinases: ERK1/2, MEK1/2, GSK3A/B, IGF1R, JNK, KRAS, MEK1/2, mTOR, PDPK1, PI3K, p38. Interestingly, in the FLT3^ITD-TKD^ model, the combined inhibition of JNK and FLT3, exclusively, *in silico* restores the TKI sensitivity, as revealed by the evaluation of the apoptosis and proliferation levels (**Fig. 4B-C**).

We thus tested *in vitro* whether the pharmacological suppression of JNK using a highly selective inhibitor could increase the sensitivity of FLT3^ITD-TKD^ cells to TKI treatment. Our data indicate that JNK plays a crucial role in cell survival of FLT3-ITD cells, since its pharmacological inhibition (SP600125) alone or in combination with Midostaurin (PKC412) significantly increased the percentage of apoptotic FLT3^ITD-TKD^ cells (**Fig. 4D**). Remarkably, the apoptosis of FLT3^ITD-TKD^ patients-derived blasts is increased upon pharmacological inhibition of JNK (**Fig, 4F**). Consistently, in these experimental conditions, we observed a significant reduction of proliferating FLT3^ITD-TKD^ cells versus cells treated with Midostaurin alone (**Fig. 4E**). Additionally, in agreement with the models’ predictions, we demonstrated that pharmacological suppression of ERK1/2 or p38 kinases have no impact on the TKI sensitivity of FLT3^ITD-TKD^ cells (**Fig. S6A-B**).

We next sought to characterize the functional role of JNK in this response. Recently, we revealed that the cell cycle controls the FLT3^ITD-TKD^ TKI resistance via the WEE1-CDK1 axis (Massacci et al., 2023). Interestingly, JNK has already been shown to play a role in cell cycle regulation through the inactivation of CDC25C, a phosphatase and positive regulator of CDK1 (Gutierrez et al., 2010). Thus, we investigated whether pharmacological inhibition of JNK may differently impact CDK1 activity in FLT3^ITD-JMD^ and FLT3^ITD-TKD^ cells. In line with our previous findings (Massacci et al., 2023; Pugliese et al., 2023), in FLT3^ITD-JMD^ cells, Midostaurin treatment increases the dephosphorylated, cytosolic, and monomeric pool of CDK1 and inactivates CDK2 (**Fig. 4G**), leading to cell accumulation in the G1 phase (**Fig. 4H**). Combined treatment of SP600125 and Midostaurin increases CDK1 and CDK2 phosphorylation and Cyclin B1 levels, increasing the percentage of G2-M and S-phases cells, compared with Midostaurin treatment alone (**Fig. 4G-H**).

As expected, in cells expressing FLT3^ITD-TKD^, Midostaurin treatment triggers the formation of the inactive stockpiled pre-M-phase Promoting Factor (MPF) (Massacci et al., 2023), constituted by the CDK1-CyclinB1 complex (**Fig. 4G, I**). This complex is associated with a significant accumulation of proliferating FLT3^ITD-TKD^ cells in the G2-M phase as compared to Midostaurin-treated FLT3^ITD-JMD^ cells. In line with these observations, CDK2 phosphorylation on activating Thr160 was significantly increased (**Fig. 4G, S6E**). On the other hand, combined treatment of SP600125 and Midostaurin induces dephosphorylation of CDK1 on the inhibitory Tyr15 and a mild accumulation of FLT3^ITD-TKD^ cells in G2-M phase (**Fig. 4G, I, S6C-D, S6G**). These observations support the hypothesis that the combination of JNK inhibition with Midostaurin treatment impacts the cell cycle progression in TKI-resistant FLT3^ITD-TKD^ cells, impairing their survival and reactivating the TKI-induced apoptosis.

### Generation of AML patient-specific logic models

Our genotype-specific Boolean models were built on *in vitro* signaling data and enabled us to formulate reliable mechanistic hypotheses underlying TKI resistance in our AML cellular models. As outlined in **Figure 5A**, to exploit their predictive power in a more clinical setting, we implemented a computational strategy that combines the models’ topological structure with patient-derived gene expression data.

**Figure 5.**
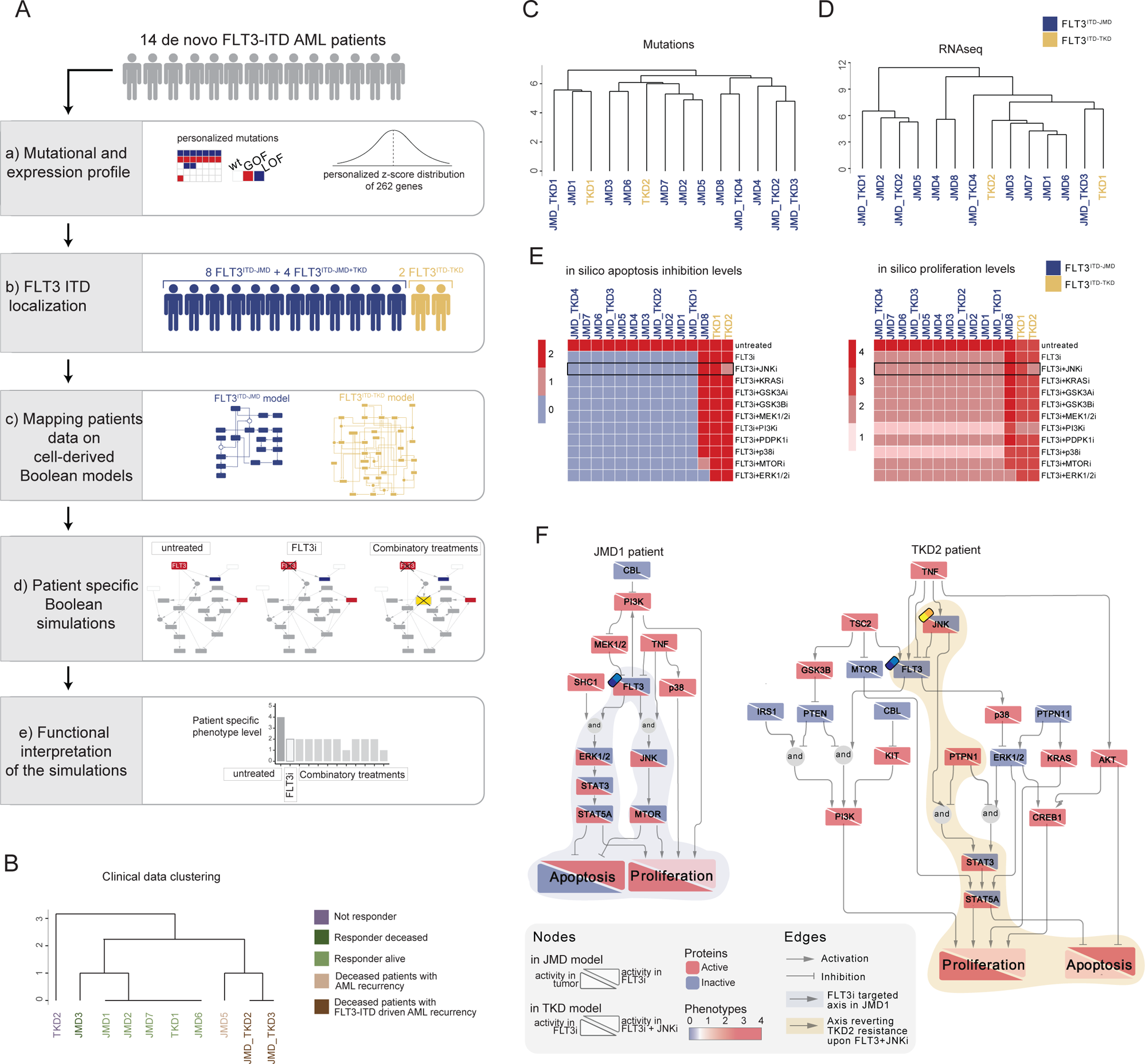
FLT3-ITD patient-specific Boolean models. **A)** Schematic representation of our computational approach to obtain personalized logic models. **B)** Hierarchical clustering of patients according to their clinical characteristics (response to chemotherapy, vital status, and AML recurrence). Resistant, alive or deceased responders, and deceased with general or FLT3-ITD AML recurrence patients are reported in purple, light or dark green, and light or dark brown, respectively. **C-D**) Hierarchical clustering of patients according to their mutational profile (**C**) and their expression profile of 262 genes (**D**). **E)** Heatmap representing patient-specific *in silico* apoptosis inhibition (*left panel)* and proliferation levels (*right panel*) upon each simulation condition. **F)** Patient-specific (JMD1 and TKD2) Boolean models. In the JMD1 model (left panel), nodes’ activity has been simulated in control (bottom-left part) and *FLT3 inhibition* conditions (upper-right part). In the TKD2 model (right panel), nodes’ activity has been simulated in *FLT3 inhibition* (bottom-left part) and *FLT3 and JNK co-inhibition* conditions (upper-right part).

As a pilot analysis, we analyzed the mutational and expression profiles of 262 genes (**Table S7**), relevant to hematological malignancies in a cohort of 14 FLT3-ITD positive *de novo* AML patients (**Fig. 5A, panel a**). Briefly, the classification of these 10 patients according to their ITD localization (see Methods) was as follows: 8 patients with FLT3^ITD-JMD^, 4 with FLT3^ITD-JMD+TKD^, and 2 with FLT3^ITD-TKD^ (**Fig. 5A, panel b**). The specific insertion sites of the ITD in the patient cohort are shown in **Table S8.** Follow-up clinical data were available for 10 out of 14 patients (**Fig. 5B, Table S9**).

Mutation profiling analysis of the patient cohort revealed a heterogeneity in the genetic background among patients and a high number of co-occurring genetic alterations (**Fig. S7A-B, Table S9**). By computing the genes’ z-scores with respect to each patient’s gene expression distribution, we detected patient-specific up- or down-regulated transcripts (**Fig. 5A, panel a, Table S9**).

Significantly, patients’ unsupervised hierarchical clustering according to the mutational profile or according to the z-score distribution of the gene expression and principal component analysis (PCA) of the gene expression data was unable to stratify patients based on their FLT3-ITD subtypes (**Fig. 5C-D, Fig. S7C**).

At this point, we tested whether we could exploit the cell-derived Boolean models to generate personalized predictive models able to reproduce the clinical outcome of patients and then identify novel personalized combinatorial treatments.

To these aims, each patient’s mutational profile (**Fig. S7A and D, Table S9**) was first used to match the suitable cell-derived FLT3-ITD model and then exploited to set the initial condition and obtain 14 personalized Boolean models (**Fig. 5A, panel c**). Next, for each patient, we performed a simulation of the following conditions *in silico*: i) *untreated* state; ii) *FLT3i* condition (see Methods); and iii) combination of FLT3i and inhibition with previously tested kinases. Importantly, our approach enabled us to obtain patient-specific predictive Boolean models able to describe the drug-induced signaling rewiring (**Fig. 5F and Fig. S8)** and to quantify ‘apoptosis inhibition’ and ‘proliferation’ levels (**Fig. 5A, panel d and e, Fig. 5E and Fig. S7E**). The anti-proliferative and pro-apoptotic response to FLT3 inhibition (**Fig. 5E**) of JMD1, JMD2, JMD3, JMD7, and JMD6 models was confirmed by follow-up clinical data that displayed a favorable outcome upon treatment (**Fig. 5B**). In fact, in JMD1 patient, the sole FLT3 inhibition impairs the STAT3-STAT5 and JNK-MTOR axes and leads to an anti-tumoral phenotype (**Fig. 5F**). Conversely, simulations of JMD5, JMD_TKD2, JMD_TKD3 and TKD1 models showed an opposite outcome with respect to real-life clinical observations (**Fig. 5B**). For example, in our *in silico* model, the clinically responder TKD1 patient (**Fig. 5B**) was resistant to all the tested combinatorial treatments, with a weak effect of PI3Ki on the pro-proliferative axis (**Fig. 5E and Fig. S7E**). One possible explanation is that the more complex mutational landscape of the TKD1 patient cannot be recapitulated by our scaffold model (**Fig. S7D**). Interestingly and in line with our previous cell-line-based findings, JNK inhibition appeared to be a promising approach to alleviate the resistant phenotype of the clinically-not-responder TKD2 (**Fig. 5B**), as revealed by the diminished levels of ‘apoptosis inhibition’ and ‘proliferation’ (**Fig. 5E-F**). Our model suggests that the effect of combinatorial FLT3i and JNKi treatment increases AML cell death through the STAT3/STA5A axis (**Fig. 5F**). Overall, this analysis showcases the two trained Boolean logic models have predictive power and can contribute to identifying potential therapeutic strategies improving clinical outcome of FLT3^ITD-TKD^ patients.

## Discussion

Cancer is primarily a signaling disease in which gene mutations and epigenetic alterations drastically impact crucial tumor pathways, leading to aberrant survival and cell proliferation. Indeed, nearly all molecularly targeted therapeutic drugs are directed against signaling molecules (Min and Lee, 2022). However, the success of targeted therapies is often limited, and drug resistance mechanisms arise, leading to therapy failure and dismal patient prognosis. To address this issue, a comprehensive, patient-specific characterization of signaling network rewiring can offer an unprecedented opportunity to identify novel promising, personalized combinatorial treatments.

Logic-based models have already been proven to successfully meet this challenge, thanks to their ability to condense the signaling features of a system and to infer the response triggered by genetic and chemical perturbations to the system *in silico* (Lee et al., 2012, 2012; Montagud et al., 2022).

In the present study, we optimized a methodology to investigate drug sensitivity using genotype-specific Boolean models. Our approach involved building a model representing the patient-specific cell state or disease status and inferring novel combinatorial anti-cancer treatments that may overcome drug resistance.

Here, we specifically applied this methodology to acute myeloid leukemia (AML) patients carrying the internal tandem duplication (ITD) in the FLT3 receptor tyrosine kinase. Our group recently showed, by integrating unbiased mass spectrometry-based phosphoproteomics with literature-derived signaling networks, that the location of the ITD insertion affects the sensitivity to TKIs therapy through a WEE1-CDK1 dependent mechanism. Our work enabled us to obtain a nearly complete, though static, picture of how FLT3-ITD mutations rewire signaling networks.

The main goal of the present study was the generation of clinically relevant, predictive models of the FLT3-ITD-dependent cell state. We aim to use these models to predict *in silico* the TKI sensitivity upon multiple simultaneous perturbations (i); to generate personalized models by combining patient-specific genomic and transcriptomic datasets (ii) and to propose novel, effective, patient-specific anti-cancer treatments (iii).

By taking advantage of the previously developed Cell Network Optimizer software (Terfve et al., 2012), we employed a multi-step strategy that trains a prior-knowledge signaling network (PKN) with a large-scale multiparametric dataset by using a Boolean logic modeling formalism.

First, we generated a literature-derived FLT3-ITD-centered signaling network encompassing relevant pathways in AML, including the regulation of key phenotypes, such as apoptosis and proliferation. Manually-curated data were made publicly available and can be freely explored by using tools offered by the SIGNOR resource website or downloaded for local analysis, in compliance with the FAIR principles (Wilkinson et al., 2016).

Second, we used the xMAP technology to interrogate signaling in FLT3-ITD cells treated with a panel of nine different perturbations. Analysis of this large multiplex signaling dataset, consisting of 16 distinct experimental conditions, revealed a clear separation between TKI-treated and untreated cells as well as IGF1 and TNFα-stimulated cells. Surprisingly, there was no clear separation between FLT3^ITD-JMD^ and FLT3^ITD-TKD^ cells, upon clustering based on signaling parameters. This may be caused by the targeted nature of our measurements, in line with our recent demonstration that unbiased phosphoproteome profiles discriminate FLT3-ITD cells according to the ITD location (JMD region vs TKD region). We also speculate that the different ITD insertion site has a less pronounced effect on cell signaling as compared to the pharmacological inhibition of key kinases (e.g., FLT3) or stimulation with cytokines. This observation highlights the necessity of a systems-based approach allowing the generation of predictive, genotype-specific models describing how signaling rewiring may affect TKI sensitivity.

Third, we optimized two genotype-specific Boolean models to delineate the signaling networks downstream of FLT3^ITD-JMD^ and FLT3^ITD-TKD^. The topology of the two Boolean models was different and most of the interactions cell-specific, suggesting a deep rewiring of the signal downstream of FLT3 due to the different locations of the ITD. Remarkably, when we simulated the pharmacological suppression of FLT3 *in silico*, our models were able to recapitulate the well-documented differential sensitivity to TKI treatment of cells expressing FLT3^ITD-JMD^ versus FLT3^ITD-TKD^. Additionally, by taking advantage of two independent publicly available datasets, including phosphoproteomic and drug sensitivity screening datasets, we validated the predictions of our models.

Simulation of several simultaneous perturbations of these models *in silico* highlighted the role of JNK in the regulation of TKI sensitivity. Remarkably, we discovered that JNK impacts the cell cycle architecture of FLT3^ITD-TKD^ cells, by acting as a mediator of the CDK1 activity. This is in line with our previously described model, showing that hitting cell cycle regulators triggers apoptosis of FLT3^ITD-TKD^ cells (Massacci et al., 2023).

In the present study, we also investigated the clinical relevance of our optimized Boolean models, in a pilot cohort of patients. By integrating the mutation and transcriptome profiling of 14 FLT3-ITD AML patients with our cell-derived logic models, we were able to derive patient-specific signaling features and enable the identification of potential tailored treatments restoring TKI resistance. To note, in our pilot analysis, we could observe that while our predictions were confirmed by follow-up clinical data for some patients (JMD1, JMD2, JMD3, JMD6, JMD7, JMD_TK2, JMD_TKD3, TKD2), the high genetic complexity of other FLT3-ITD positive patients was not completely addressed by our cell line-derived scaffold model (JMD5, TKD1). This could be due to a number of factors: i) the size of the FLT3-ITD patient subgroups may have been too small to derive significant biological conclusions (e.g., only two patients with FLT3^ITD-TKD^); ii) the panel of molecular readouts in our training dataset might be too limited to capture the pleiotropic impact of the FLT3-ITD mutations; and iii) a more heterogeneous experimental data might be needed to train a predictive model able to recapitulate the genetic background of a real cohort of patients.

In conclusion, the integration of a cell-based multiparametric dataset with a prior knowledge network in the framework of the Boolean formalism enabled us to generate optimized mechanistic models of FLT3-ITD resistance in AML. This is the proof of concept that our personalized informatics approach described here is clinically valid and will enable us to propose novel patient-centered targeted drug solutions. In principle, the generalization of our strategy will enable us to obtain a systemic perspective of signaling rewiring in different cancer types, driving novel personalized approaches.

## Abbreviations

FLT3: Fms Related Receptor Tyrosine Kinase 3
CDK1: Cyclin-dependent kinase 1
CDK2: Cyclin-dependent kinase 2
TKI: tyrosine kinase inhibitor
ITD: internal tandem duplication
JMD: juxtamembrane domain
TKD: tyrosine kinase domain
PKN: prior knowledge network
KINi: Kinase inhibitors

## Acknowledgments

We thank Prof. Gianni Cesareni and Prof. Luisa Castagnoli for their essential scientific input. Dr. Serena Paoluzi for their technical support. This research was funded by the Italian association for cancer research (AIRC) with a grant to F.S. (Start-Up Grant n. 21815). S.L., V.V., and S.G. are supported by MUR, G.M.P., G.M., and V.B. are supported by the AIRC Start-up grant number 21815.

## Author contributions

Conceptualization, S.L., V.V., L.P., F.S.; methodology, S.L., V.V., V.B., G.M., L.P., F.S.; formal analysis, S.L., V.V., G.M.P., S.G., V.B., G.M., P.C., J.D.M., G.P., G.M., M.I.; investigation, S.L., V.V., with the contribution of T. F., D. M., M. B., M.H.C.; writing original draft preparation, S.L., V.V., F.S., L.P.; resources, F.S.; writing review and editing, all; supervision, L.P., F.S.; funding acquisition, F.S. All authors have read and agreed to the published version of the manuscript.

## Materials and Methods

### Cell culture

Murine Ba/F3 cells stabling expressing ITD-JMD and ITD-TKD constructs were kindly provided by Prof. T. Fischer. Cells were cultured in RPMI 1640 medium (Hyclone, Thermo Scientific, Waltham, MA) supplemented with 10% heat-inactivated fetal bovine serum (ECS0090D Euroclone, Italy, MI), 100 U/ml penicillin and 100 mg/ml streptomycin (Gibco 15140122), 1 mM sodium pyruvate (Sigma-Aldrich, St. Louis, Missouri, United States, S8636) and 10 mM 4-(2-hydroxyethyl)-1-piperazineethanesulfonic acid (HEPES) (Sigma H0887). These cells were chosen as an experimental system as previously described (Massacci et al., 2023).

### Multiparametric experiment of signaling perturbation

Ba/F3 FLT3^ITD-JMD^ and FLT3^ITD-TKD^ cells were cultured in complemented RPMI w/o FBS for 16 hours. Afterward, cells were treated with a panel of small molecule inhibitors for 90 minutes: Midostaurin 100nM (Selleck chemical, S8064), SB203580 10μM (Selleck chemical, S1076), SP600125 10μM (Selleck chemical, S1460), Wortmannin 35nM (Selleck chemical, S2758), Rapamycin 100nM (Sigma-Aldrich, R8781), UO126 15μM (Sigma-Aldrich, 662005), LY2090314 20nM (Selleck chemical, S7063). Cells were stimulated with IGF1 100ng/ml (Sigma-Aldrich, I8779), and TNFα 10ng/ml (Miltenyi Biotec, 130-101-687). The table below summarizes the treatments used, inhibitors and stimuli, their specific targets, the readout sentinels, and the concentrations and the treatment time chosen. We selected the inhibitors for their specificity towards key kinases in the FLT3-ITD downstream signaling. We tested their efficacy in our model cell line to set the optimal concentration and time to inhibit the kinase activity to phosphorylate its downstream targets (i.e. the UO126 at 15uM for 90 minutes inhibits MEK and we observed de-phosphorylated ERK).

We selected IGF1 and TNFα as stimuli to fully activate the receptors and their downstream kinases in order to perturb and measure more efficiently the signaling. Each treatment and perturbed kinase were paired with a sentinel analyte to monitor the responses to perturbations of all main signal transduction pathways of our cell lines included in the PKN.

We combined the treatments listed in **Table 2** to finally obtain 16 different experimental conditions in both FLT3^ITD-JMD^ and FLT3^ITD-TKD^ cell lines. The experimental conditions are summarized in **Table S2** and listed below:

1. FLT3i
2. FLT3i+IGF1
3. FLT3i+TNFα
4. FLT3i+p38i+ TNFα
5. FLT3i+JNKi+ TNFα
6. FLT3i+PI3Ki+IGF1
7. FLT3i+mTORi+IGF1
8. FLT3i+MEKi+IGF1
9. IGF1
10. TNFα
11. p38i+ TNFα
12. JNKi+ TNFα
13. PI3Ki+IGF1
14. mTORi+IGF1
15. MEKi+IGF1
16. GSK3i+IGF1

**Table 2.** Small molecule inhibitors and stimuli for the multiparametric analysis.

| inhibitors | target | usage | time | stimuli | usage | time |
| --- | --- | --- | --- | --- | --- | --- |
| Midostaurin | FLT3 | 100nM | 90 minutes | IGF1 | 100ng/ml | 10 minutes |
| SB203580 | p38 | 10μM | 90 minutes | TNFα | 10ng/ml | 10 minutes |
| SP600125 | JNK | 20μM | 90 minutes |  |  |  |
| Wortmannin | PI3K | 50nM | 90 minutes |  |  |  |
| Rapamycin | mTOR | 100nM | 90 minutes |  |  |  |
| UO126 | MEK1/2 | 15μM | 90 minutes |  |  |  |
| LY2090314 | GSK3 | 20nM | 90 minutes |  |  |  |

We combined the inhibition of a specific target with the stimulation of corresponding pathways with either IGF1 (AKT-MAPK pathway) or TNFα (p38-JNK pathway), in the presence or absence of FLT3 inhibitor Midostaurin. Combinatorial treatments aimed at perturbing the cell signaling and at measuring amplified signaling changes in our system. We therefore measured the phosphorylation levels of 14 sentinel proteins listed in **Table 3** through the X-Map Luminex technology. To each residue measure, we mapped the functional role associated, activatory =1, inhibitory =-1 depending on the annotated function on PhosphoSitePlus.

**Table 3.** xMAP analytes.

| analytes | cat.no. | phosphosite measured | activity annotation |
| --- | --- | --- | --- |
| CREB1 | 42-680MAG | Ser133 | 1 |
| ERK1/2 | 42-680MAG | Thr185/Tyr187 | 1 |
| JNK | 42-680MAG | Thr183/Tyr 185 | 1 |
| p38 | 42-680MAG | Thr180/Tyr182 | 1 |
| STAT3 | 42-680MAG | Ser727 | 1 |
| STAT5 | 42-680MAG | Tyr694/699 | 1 |
| p70S6K | 42-611MAG | Thr412 | 1 |
| RPS6 | 42-611MAG | Ser235/236 | 1 |
| MTOR | 42-611MAG | Ser2448 | 1 |
| IGF1R | 42-611MAG | Tyr1135/Tyr1136 | 1 |
| PTEN | 42-611MAG | Ser380 | -1 |
| TSC2 | 42-611MAG | Ser939 | -1 |
| GSK3A | 42-611MAG | Ser21 | -1 |
| GSK3B | 42-611MAG | Ser9 | -1 |
| B-Tubulin | 46-413MAG | total protein | loading control |

Cells were collected, lysed, and stained following the manufacturer’s instructions. Briefly, in 96 well plates, cell lysates were marked with the mix of specific antibodies covalently bound to magnetic beads, and the signal was amplified with a biotin-streptavidin system. The plates were read through the Magpix instrument: for each sample, the instrument measured the intensity of the fluorescent signal pairing it with the identity of the beads given by their location on the magnetic field. As the final output, we obtained the median fluorescence intensity (MFI) for all the sentinels in each experimental condition, paired with the number of detected beads. For each sentinel, the fluorescent threshold should be associated with a count of more than 50 beads to be technically reliable. We then, excluded from the dataset the measures with less than 50 beads detected as shown in the “filter on n. beads” sheet of **Table 3,4**. Then, we normalized the MFI of each analyte on the values of B-Tubulin as loading control and we calculated the median and SD of the three biological replicates (**Table S3,4**).

### Data normalization

The phosphorylation measure of the 14 sentinel signaling proteins was scaled between 0 and 1, using customized Hill functions for each analyte. By applying the formula:

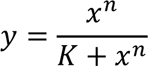

We derived n and K parameters of customized Hill functions from the distribution of each analyte in our experimental data. Briefly, given the asymptotic behavior of the Hill function, we set the experimental maximum (maxS) of the analyte to be 0.999 (theoretical maximum, maxT) and the experimental minimum (minS) to be 0.001 (theoretical minimum, minT). We then computed the b parameter as:

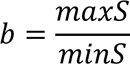

Next, we calculated n and K for each analyte Hill function:

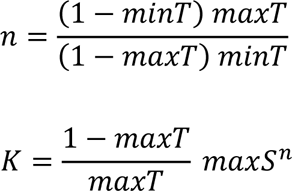

### Principal Component Analysis and Hierarchical Clustering

Principal component analysis was performed using the *stats* R package (v. 4.1.2).

Perseus software was employed to perform unsupervised hierarchical clustering. Specifically, the Pearson correlation between phosphorylation profiles of sentinel proteins across different experimental conditions was calculated and used to generate the three. Similar experimental conditions are in the same branches.

### FLT3 ITD-specific Boolean model construction with CellNetOptimizer

We exploited the CellNetOptimizer pipeline which integrates (i) a prior knowledge network (PKN) and (ii) multi-parametric, normalized experimental data to obtain two FLT3 ^TD-JMD^ and ^-TKD^ dynamic and predictive Boolean models. An extended and detailed step-by-step description of the whole modeling strategy is available in Supplementary Material and the code to reproduce the analysis is available at https://github.com/SaccoPerfettoLab/FLT3-ITD_driven_AML_Boolean_models.

### Prior Knowledge Network manual curation

We built a FLT3-ITD specific prior knowledge network (PKN) combining (i) a manual curation and (ii) a data-driven approach. Starting from the SIGNOR database, through a curation effort, we assembled a causal network describing the FLT3-ITD signaling, comprising all the direct and indirect interactions implied in the receptor signaling and leukemogenesis. The PKN is publicly available on SIGNOR (https://signor.uniroma2.it/pathway_browser.php?organism=&pathway_list=SIGNOR-Sara). We downloaded the interaction table, and we manually simplified the network, we compressed some articulated and redundant paths (**Table S1**). We converted the network into a .sif file made of three columns, entity A, entity B, and interaction type described with 1 if activatory or −1 if inhibitory. Importantly, during the optimization process, the PKN was adjusted until we reached an optimal performance of the model. The final version of the PKN displayed 76 nodes and 193 edges. Next, we used CellNOptR v.1.40.0, to preprocess the PKN and to convert the causal network into logical functions (scaffold model), describing the regulatory relations among gene products using OR/AND Boolean operators. Briefly, we first compressed and expanded the PKN (Terfve et al., 2012) to obtain a network of 204 nodes (30 proteins and 176/178 AND Boolean operators) and 612 edges. We, next, exploited CNORfeeder v.1.34.0 to impute missing links derived directly from the experimental data, using the FEED method, developed specifically to infer signaling networks from perturbation experiments. Finally, we integrated the two networks obtaining a scaffold model of 756 edges for FLT3^ITD-JMD^ (144 data-driven) and 782 edges for ^ITD-TKD^ (170 data-driven).

### Network model optimization

We performed 1000 runs of optimization using CellNOptR which creates context-specific Boolean models (i) by filtering out interactions not relevant to the system and (ii) by selecting the Boolean operators (i.e., AND/OR) that best integrate inputs acting on the same node.

CellNOptR exploits a genetic algorithm that minimizes the difference (mean squared error, mse) between experimental data and the values simulated from the Boolean model.

We sorted the models according to mse in ascending order and selected the first 100 models (family of best models). Then, we calculated the average state of each protein in the 100 best models. This procedure enables quantitative prediction even using Boolean models, which are discrete by nature. These averaged values were compared with the training data to evaluate the goodness of fit. We used the model with the lowest mse of FLT3^ITD-JMD^ and TKD cell lines for further analyses (best model). These final models accounted for 68 and 60 nodes (of which 38 and 30 are AND operators) and 161 and 133 edges for FLT3^ITD-JMD^ and FLT3^ITD-TKD^, respectively. To keep a measure of the whole optimization procedure in the best models, we added as edges’ attribute the frequency of each edge in the family of 100 models and we considered as ‘high confidence edges’ the ones having a frequency of 0.4 (edges present in the final model of 40 stochastic optimization procedures out of 100).

### Boolean models’ validation using independent resources

Using the simulatorT1 function of CellNOptR, we computed the steady state of FLT3^ITD-JMD^ and ^ITD-TKD^ Boolean models with and without the inhibition of FLT3 and other druggable nodes. To independently validate the models, we used as a reference the phosphoproteomic data of FLT3-ITD Ba/F3 cells (FLT3^ITD-JMD^ and FLT3^ITD-TKD^) upon quizartinib (AC220) treatment (Massacci et al., 2023). We mapped xMAP residues associated with protein complexes (e.g. ERK1/2) to unique protein sequences (e.g., Mapk1 and Mapk3) (**Table S2**). We estimated the activity of sentinel proteins in the reference dataset using the modulation of their regulatory phosphosites. Then, we compared the estimated activity with the sentinels’ states in the FLT3 inhibition simulation.

Moreover, to functionally interpret the results and assess the reliability of the model, we computed the activity of ‘apoptosis’ and ‘proliferation’ phenotypes upon FLT3 and other druggable nodes inhibition after the annotation of model proteins as pro- and anti-apoptotic (or proliferative). To obtain the table of protein annotations, with proximal phenotypes, we exploited a recently in-house developed method, dubbed ProxPath, (Iannuccelli et al., 2023) which computes significantly ‘close’ paths linking SIGNOR proteins and phenotypes. The distance table connecting the model nodes to the ‘Apoptosis’ and ‘Proliferation’ phenotypes is available in **Table S1.** To compute the phenotypes’ activation status, we integrated with the OR logic (‘sum of scores’) the activation status of upstream nodes, which were also endpoint proteins in high-confidence signaling axes (edge frequency 0.4) in the cell-specific models. As such, if two regulators of the same phenotype were linked in the same axis, we considered only the one at the end of the cascade.

The Beat AML program on a cohort of 672 tumor specimens collected from 562 patients has been exploited for model validation. We focused on drug sensitivity screening on 134 patients carrying the typical FLT3-ITD mutation in the JMD region. Drugs were annotated for their targets using SIGNOR and ChEBI databases. Drugs inhibiting FLT3, PI3K, mTOR, JNK, and p38 were selected and the average IC50 of FLT3^ITD-JMD^ patient-derived primary blasts was calculated (**Table S6**).

### Combinatory treatment inference

To identify promising cotreatments able to revert the resistant phenotype, we exploited the predictive power of the generated Boolean models and performed an *in silico* knock-out of key kinases present in the FLT3^ITD-TKD^ model (ERK1/2, MEK1/2, GSK3A/B, IGF1R, JNK, KRAS, MEK1/2, MTOR, PDPK1, PI3K, p38). Briefly, we computed the steady state of each cell line best model before and after the co-inhibition of FLT3 and every key kinase (11 possible combinations). Then we inferred the activity of ‘apoptosis’ and ‘proliferation’ phenotypes. We eventually selected co-treatments in the FLT3^ITD-TKD^ model able to trigger activation levels of the ‘apoptosis’ and ‘proliferation’ to the same level as the FLT3^ITD-JMD^ model.

### Apoptosis assay

Ba/F3 cells were treated with Midostaurin 100nM and SP600125 10μM for 24 hours.

The concentration of SP600125 to use for this long-term treatment was chosen based on setup experiment: we treated sensitive and resistant cells with increasing concentrations of SP600125 for 24 hours and evaluated the cell viability using the Cell Proliferation Kit I (MTT) (Roche, Cat. 11465007001) and measuring the absorbance value at λ=590nm. We then calculated the IC50 with a nonlinear regression drug-response curve fit using Prism 7 (GraphPad). IC50 values are approximately 1.5 µM in FLT3-ITD mutant cell lines (FLT3^ITD-JMD^ cells IC50=1.54μM; FLT3^ITD-TKD^ cells IC50=1.69μM). The SP600125 treatment affects cell viability, reaching a plateau phase of cell death and at about 2 µM. I used 10µM SP600125 to inhibit JNK phosphorylation. (Kim et al., 2010; Moon et al., 2009). The concentration of Midostaurin for the apoptotic assay was chosen based on the previously published work (Massacci et al., 2022) where we show that FLT3^ITD-TKD^ cells treated with Midostaurin 100nM show lower apoptotic rate and higher cell viability compared to FLT3^ITD-JMD^ cells. Apoptotic cells were analyzed with the Ebioscience™ Annexin V Apoptosis Detection Kit APC according to the kit instructions (Cat. 88-8007-74, Thermo Fisher Scientific. Samples were read through the CytoFLEX S (Beckman Coulter) instrument using the APC laser to detect the Annexin-V+ cells. Quality control of the cytometer was assessed weekly using CytoFLEX Daily QC Fluorospheres (Beckman Coulter B53230). The fluorescence threshold was set for the APC laser using a blank sample, without the fluorescent label. The results were analyzed by the CytExpert software and represented in bar plots as the percentage of Annexin-V+ cells fold change of treated conditions on controls.

### MTT assays

Ba/F3 cells were treated with Midostaurin 100nM, SP600125 20μM, SB203580 10μM, and UO126 15μM for 24 hours in 96-well plates. The concentration of SB203580 and UO126 to use for this long-term treatment was chosen based on previous data available in the lab and set-up experiments. We used the concentration in which we could observe a reduced activity of the target (lower phosphorylation of downstream proteins) without cell toxicity. Cell viability was assessed using the Cell Proliferation Kit I (MTT) (Roche, Cat. 11465007001), following the manufacturer’s instructions. The plates were read through a microplate reader (Bio-Rad) at λ=590 nm. The results were represented in bar plots as fold change of treated conditions on controls.

### Cell cycle analysis

Ba/F3 cells were treated with Midostaurin 100nM and SP600125 10μM for 24 hours, 106 cells were collected, washed in ice-cold PBS 1X, and fixed in agitation with 70% cold ethanol. Fixed samples were incubated at 4°C O/N, washed in PBS 1X, and resuspended in 1μg/ml DAPI (Thermo Scientific, #62248) and 0.2 mg/ml RNase (Thermo Scientific, # 12091021) PBS solution before analysis. Samples were read through the CytoFLEX S (Beckman Coulter) instrument using the PB450 laser to detect the DAPI+ cells. Data were analyzed by CytExpert (Beckman Coulter) software and represented in bar plots as a percentage of DAPI+ cells.

### Immunoblot analysis

Ba/F3 cells were seeded at the concentration of 5×105cells/ml and treated with Midostaurin 100nM, SP600125 20μM, SB203580 10μM, UO126 15μM, for 24 hours, and TNFα 10ng/ml for 10 minutes. Cells were lysed for 30 minutes in ice-cold lysis buffer (150 mM NaCl,50 mM Tris-HCl pH 7.5, 1% Nonidet P-40 (NP-40), 1 mM EGTA,5 mM MgCl2, 0.1% SDS) supplemented with 1 mM PMSF,1 mM orthovanadate, 1 mM NaF, protease inhibitor mixture1X, inhibitor phosphatase mixture II 1X, and inhibitor phosphatase mixture III 1X. The insoluble material was separated at 13,000 rpm for 30 minutes at 4 °C and total protein concentration was measured on supernatants using Bradford reagent (Bio-Rad). Protein extracts were denatured with NuPAGE®LDS (Invitrogen) and boiled at 95 °C for 10 minutes. SDS-PAGE and transfer were performed on 4–15%Bio-Rad Mini/Midi PROTEAN®TGX™ and Trans-Blot®Turbo™ mini/midi nitrocellulose membranes using the Trans-Blot®Turbo™ Transfer System (Bio-Rad). Nonspecific binding sites were blocked using 5% non-fat dried milk in TBS-0.1% Tween-20 (TBS-T) for 1 h at RT, shaking. Primary antibodies were diluted according to the manufacturer’s instruction and incubated at 4 °C O/N, shaking. HRP-conjugated secondary antibodies were diluted 1:3000 in 5% non-fat dried milk in TBS-T and incubated for 1h at RT, shaking. Peroxidase chemiluminescence reaction was enhanced with Clarity™Western ECL Blotting Substrates (Bio-Rad) and detected through the Chemidoc detection system (Bio-Rad). Densitometric quantitation of raw images was obtained with Fiji software (Image J, NIH). The primary and secondary used antibodies are listed: CDK1 (sc-53219); CDK2 (sc-6248); phpspho-CDK1 Y15 (CS 9111); phospho-CDK1 T161(CS 9114); phospho-CDK2 T160 (CS 2561); phospho-SAPK/JNK T183/Y185 (CS 9251); SAPK/JNK (CS 9252); Cyclin B1 (CS 4138); CyclinE2 (CS 4132); Vinculin (CS 13901).

### Primary patient blast sensitivity

Peripheral blood (PB) samples were collected from patients with acute myeloid leukemia (AML) in accordance with the Declaration of Helsinki (ethics committee approval number 115/08) and with the patients’ informed consent. The FLT3-ITD mutation integration site was determined as previously described (Massacci et al., 2023).

Mononuclear cells were isolated from PB using Ficoll-Paque (GE Healthcare, Chicago, IL). Cryoconserved peripheral blood mononuclear cells (PBMCs) from seven patients were cultured in RPMI-1640 medium (Sigma-Aldrich, St. Louis, MO) supplemented with 10% fetal calf serum (FCS) (Bio&Sell GmbH, Germany), 2 mM L-glutamine (Sigma-Aldrich), and 40 U/mL penicillin-streptomycin (ThermoFisher Scientific) at a density of 5×105/mL. Cultures were incubated for 48 hours in the absence or presence of 100 nM PKC412 and 10 µM SP600125 (all Selleck Chemicals LLC, Houston, TX), or combinations of PKC412 with SP600125. The viability of the patient’s blast cells was assessed by flow cytometry, while the specific cell death was calculated as described previously (Pugliese et al., 2023). Briefly, samples were stained with fluorochrome-conjugated antibodies against CD45, CD33, CD34, CD13, and CD117, followed by the addition of Annexin V and 7AAD. The samples were recorded using a NorthernLight-3000 spectral flow cytometer (Cytek Biosciences, Freemont, CA) and analyzed through FlowJo V10.9 (BD Bioscience, Franklin Lakes, NJ).

### Statistical analysis

Data are represented as the mean ± SEM of at least three independent experimental samples (n=3). Comparisons between three or more groups were performed using the ANOVA test. Statistical significance between the two groups was estimated using Student’s t-test. Statistical significance is defined as p value where *p< 0.05; **p<0.01; ***p< 0.001; ****p<0.0001. All statistical analyses were performed using Prism 7 (GraphPad).

### RNAseq of patient samples

RNA was extracted from peripheral blasts of 14 treatment naïve patients with *de novo* FLT3 ITD driven-AML diagnosis. Clinical characteristics were available for only 10 patients (**Table S9**). Peripheral blood (PB) samples from 14 AML patients were obtained upon the patient’s informed consent. The integration site of the FLT3-ITD mutation was determined as previously described (Rücker et al., 2022). Briefly, RNA was prepared from PBMCs using the RNeasy Mini Kit (Qiagen, Germany), retro-transcribed in cDNA, and quantified using the Qubit 4.0 fluorimetric Assay (Thermo Fisher Scientific) and sample integrity, based on the DIN (DNA integrity number), was assessed using a Genomic DNA ScreenTape assay on TapeStation 4200 (Agilent Technologies).

High throughput sequencing was performed on the coding region of 262 genes involved in hematologic malignancies. A comprehensive list of all genes is included in **Table S7**.

Genomic DNA Libraries were prepared from 100 ng of total DNA using the NEGEDIA Cancer Haemo Exome sequencing service (Next Generation Diagnostic srl) which included library preparation, target enrichment using Cancer Haemo probe set, quality assessment, and sequencing on a NovaSeq 6000 sequencing system using a paired-end, 2×150 cycle strategy (Illumina Inc.). Paired-end reads were produced with a 100x coverage, and a median of 40 M reads per sample.

### Bioinformatic analysis of transcriptome data of patient samples

The raw data were analyzed by Next Generation Diagnostics srl Cancer haemo Exome pipeline (v1.0) which involves a cleaning step by UMI removal, quality filtering and trimming, alignment to the reference genome, removal of duplicate reads and variant calling (FASTQC: https://www.bioinformatics.babraham.ac.uk/projects/fastqc/, (Freed et al., 2017; McLaren et al., 2016; Smith et al., 2017)).

Illumina NovaSeq 6000 base call (BCL) files were converted into fastq files through bcl2fastq (2019) (version v2.20.0.422). UMI removal was carried out with UMI-tools 1.1.1. Data quality control was performed with FastQC v.0.11.9 and reads were trimmed and cleaned using Trimmomatic 0.38.

Alignment to human reference (hg38, GCA_000001405.15), deduplication, and variant calling were performed with Sentieon 202011.01.

Variant calling output was converted into vcf and MAF format. The maf files were further annotated with the OncoKB™ annotator (Chakravarty et al., 2016) to obtain the mutation effect on protein function and oncogenicity. We filtered out variants with MUTATION_EFFECT equal to ‘Unknown’, ‘Inconclusive’, ‘Likely Neutral’, and ‘Switch-of-function’ (**Table S9**). All downstream analyses were carried out using R v.4.1.2 and BioConductor v.3.13 ((Huber et al., 2015); R Core Team, 2020)).

Gene counts tables were generated from bam files using Rsubread v. 2.8.2. Data were normalized with the “trimmed mean of M values” (TMM) method of edgeR v3.36.0 (Robinson et al., 2010) and converted to log2 (**Table S9**).

### FLT3 ITD localization

Generic variant callers can’t identify medium-sized insertions, like FLT3 ITDs. As such, we used the specialized algorithm getITD v.1.5.16 (Blätte et al., 2019) to localize ITDs in each patient. Reads mapping on FLT3 genomic region between JMD and TKD domain (28033888 – 28034214) were extracted from bam files and converted in fastq format with samtools v.1.16.1.

getITD was run with default parameters, but with a custom reference sequence without introns (‘caatttaggtatgaaagccagctacagatggtacaggtgaccggctcctcagataatgagtacttctacgttgatttcagagaatatgaatatgatctcaaatgggagtttccaagagaaaatttagagtttgggaaggtactaggatcaggtgcttttggaaaagtgatgaacgcaacagcttatggaattagcaaaacaggagtctcaatccaggttgccgtcaaaatgctgaaag’) that was annotated according to getITD annotation file. Patients carried multiples ITDs and we classified as follows: 8 patients as FLT3^ITD-JMD^ (JMD1-8), 4 patients as FLT3^ITD-JMD+TKD^ (JMD_TKD1-4), and 2 as FLT3^ITD-TKD^ (TKD1-2) (**Table S8**).

### Patients’ simulation on cell-derived ITD-specific Boolean models

We decided to use the FLT3^ITD-JMD^ cell models for both FLT3^ITD-JMD^ and FLT3^ITD-JMD+TKD^ given the dominant effect that the FLT3^ITD-JMD^ mutation displays over the FLT3^ITD-TKD^ one (Rücker et al., 2022).

We used patients’ data to perform Boolean simulations on cell-derived ITD-specific Boolean models. We used FLT3^ITD-JMD^ cell lines Boolean models for FLT3^ITD-JMD^ and FLT3^ITD-JMD+TKD^ patients and FLT3^ITD-TKD^ models for FLT3^ITD-TKD^ patients. Each patient mutational profile was binarized, setting “Loss-of-function” or “Likely loss-of-function” equal to 0, and “Gain-of-function” or “Likely gain-of-function” equal to 1. Using CellNOptR, for each patient we computed the steady state of its correspondent cell-derived Boolean model in two conditions: i) the input is the mutational profile (tumor simulation - “untreated”); ii) the input is the mutational profile + FLT3 inhibition (treatment with FLT3i simulation). Hence, we inferred the ‘apoptosis’ and ‘proliferation’ phenotype scores, as stated above.

We also performed an *in silico* combined treatment of patients simulating their mutations with the inhibition of FLT3 and a key signaling kinase (eleven different simulations).

### Network visualization

To visualize the best model, we exported CellNOptR results using writeScaffold and writeNetwork functions and imported the network in Cytoscape v.3.9.0. For nodes and edges’ color, we used the style of Cytocopter app v.3.9. We added as edges’ attribute the frequency of each edge in the family of 100 models and we kept only the edges having a frequency > 0.4.

### Statistical analyses and tools

The patient z-score was computed by subtracting the mean of patient-specific expression distribution and dividing by its standard deviation. PCA and clustering graphs were generated with stats v.4.1.2, ggfortify v.0.4.15, and ggdendrogram v.0.1.23 packages. Heatmaps were created using pheatmap v.1.0.12. Phenotypes bar plots are created with ggplot2 v.3.4.0.

### Data and code availability

Curated data have been submitted to SIGNOR for reuse and interoperability and can be accessed at https://signor.uniroma2.it/downloads.php. Transcriptomic data of patients has been submitted to GEO and can be accessed using the accession number: GSE247483. The code developed for the study has been organized on a GitHub page and is available at https://github.com/SaccoPerfettoLab/FLT3-ITD_driven_AML_Boolean_models.

