## Supplementary Material and Methods for "Unveiling the signaling network of FLT3-ITD AML improves drug sensitivity prediction"

### Supplementary material for Latini and Venafra et al.

#### Supplementary Figures

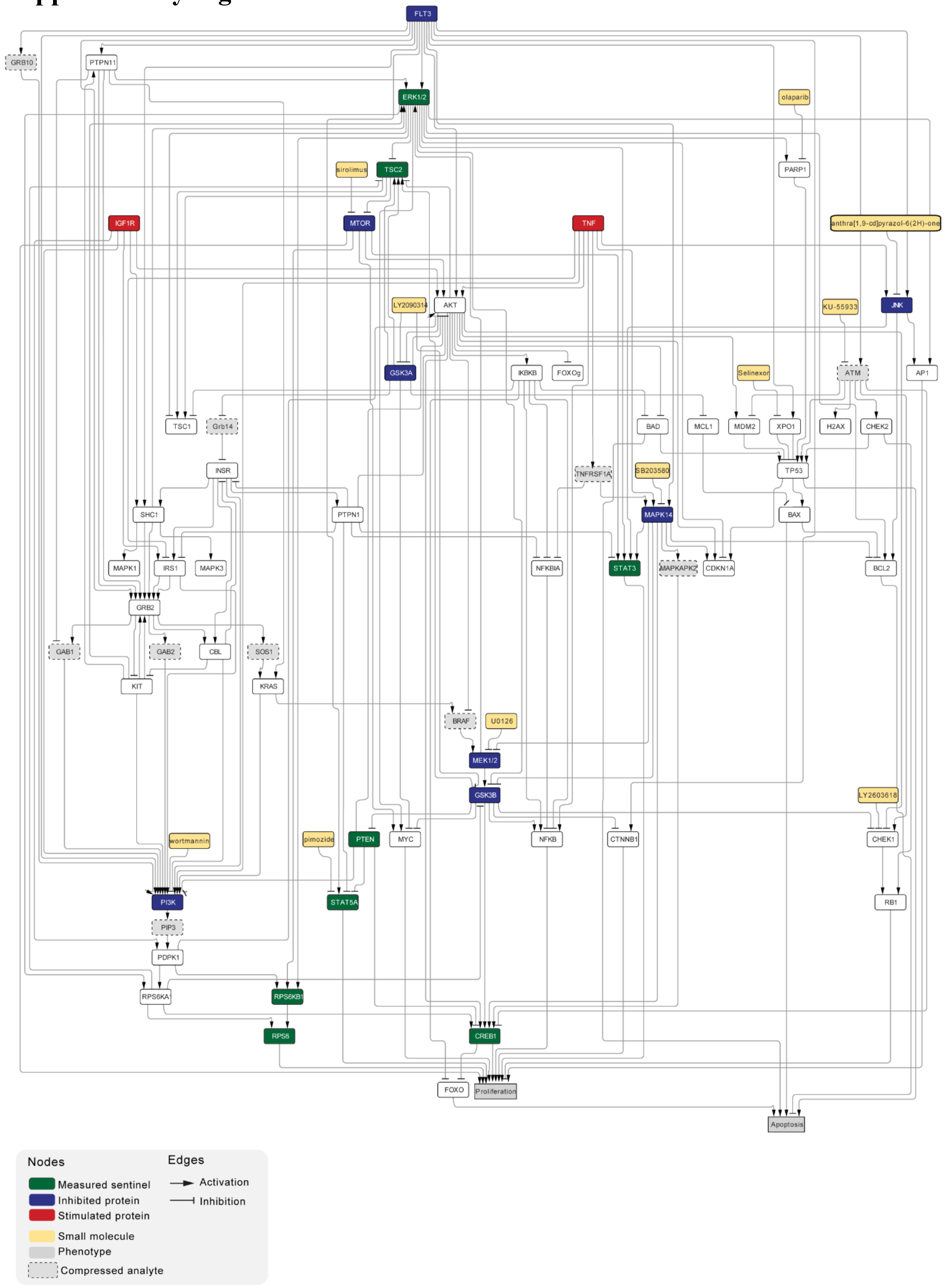

**Figure S1. FLT3-ITD manually curated Prior Knowledge Network.** Network representation of the PKN, each node represents a protein, a cytokine (green) or a small molecule inhibitor (yellow). The proteins are colored following the CNO pipeline graphics: target proteins in red, sentinel proteins in blue and essential nodes in white. The edges represent the directed interactions between the nodes: arrowhead when the interaction is activatory, hammerhead when the interaction is inhibitory.

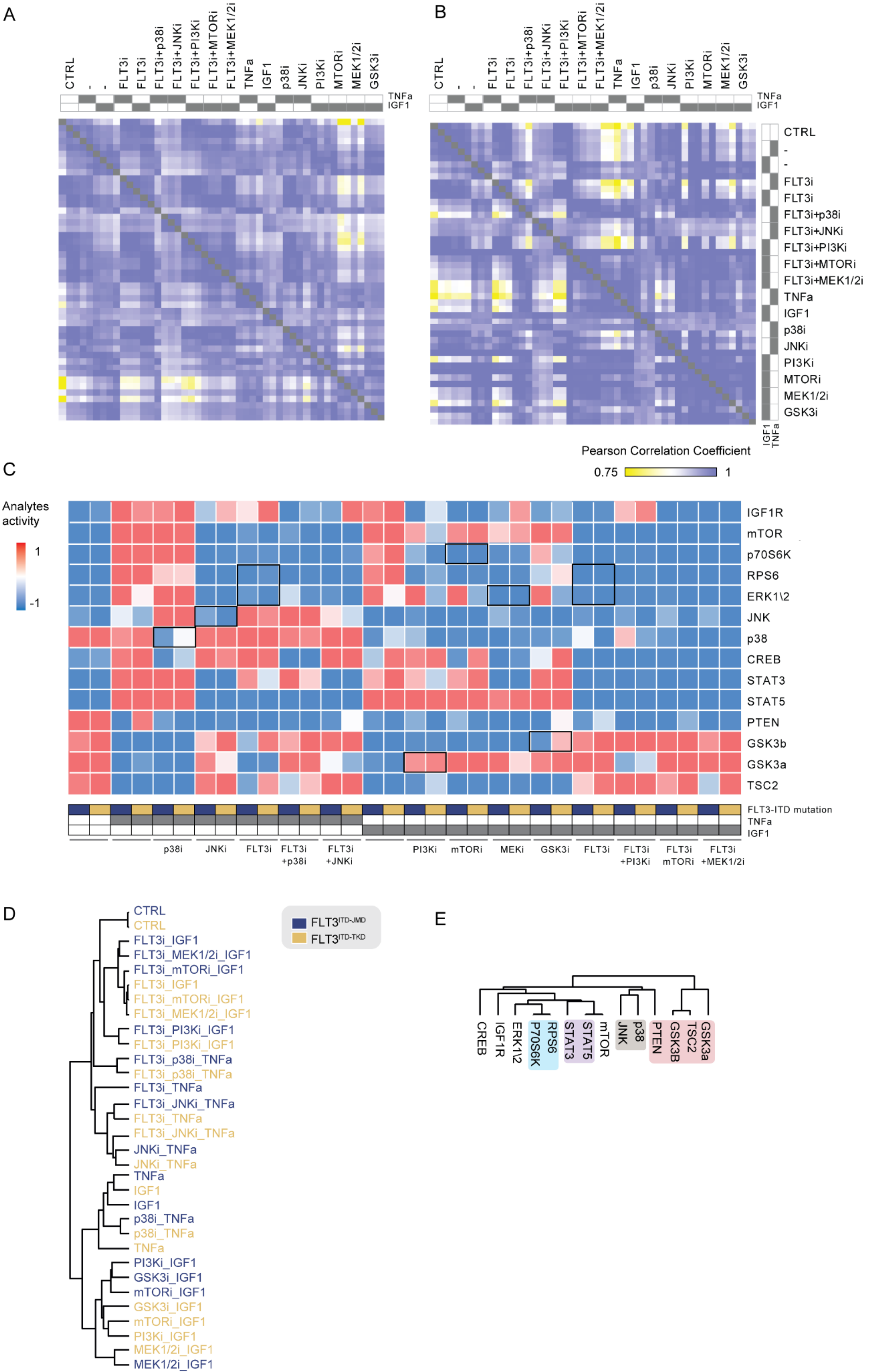
A
B
C
D
E
CTRL
-
FLT3i
FLT3i+p38i
FLT3i+JNKi
FLT3i+PI3Ki
FLT3i+MTORi
FLT3i+MEK1/2i
TNFa
IGF1
p38i
JNKi
PI3Ki
MTORi
MEK1/2i
GSK3i
Pearson Correlation Coefficient
0.75
1
Analytes activity
IGF1R
mTOR
p70S6K
RPS6
ERK1\2
JNK
p38
CREB
STAT3
STAT5
PTEN
GSK3b
GSK3a
TSC2
FLT3-ITD mutation
FLT3i +p38i
FLT3i +JNKi
mTORi
MEKi
GSK3i
FLT3i +PI3Ki
FLT3i mTORi
FLT3i +MEK1/2i
FLT3ITD-JMD
FLT3ITD-TKD
FLT3i_IGF1
FLT3i_MEK1/2i_IGF1
FLT3i_mTORi_IGF1
FLT3i_PI3Ki_IGF1
FLT3i_p38i_TNFa
FLT3i_TNFa
FLT3i_JNKi_TNFa
JNKi_TNFa
p38i_TNFa
PI3Ki_IGF1
GSK3i_IGF1
mTORi_IGF1
MEK1/2i_IGF1
CREB
P70S6K
STAT5
GSK3B

**Figure S2. Global overview of multiparametric data.**

**A-B)** Heatmap showing the Pearson correlation coefficients between the different biological replicates in FLT3 $^{ITD-JMD}$ cells (**A**) and FLT3 $^{ITD-TKD}$ cells (**B**).

**C**) Heatmap of the complete dataset representing the activity of each analyte (rows) in all the experimental conditions (columns). Each square is colored according to the activity of the analyte in each experimental condition (from active(1)=red to inactive(0)=blue). Black squares highlight the activity of specific analytes upon their direct or upstream inhibition.

**D**) Unsupervised, hierarchical clustering of the intensity of the measured sentinel proteins discriminates samples according to the treatment.

**E**) Unsupervised, hierarchical clustering allows the classification of the sentinel proteins according to their role (red) and pathway (grey, purple and blue).

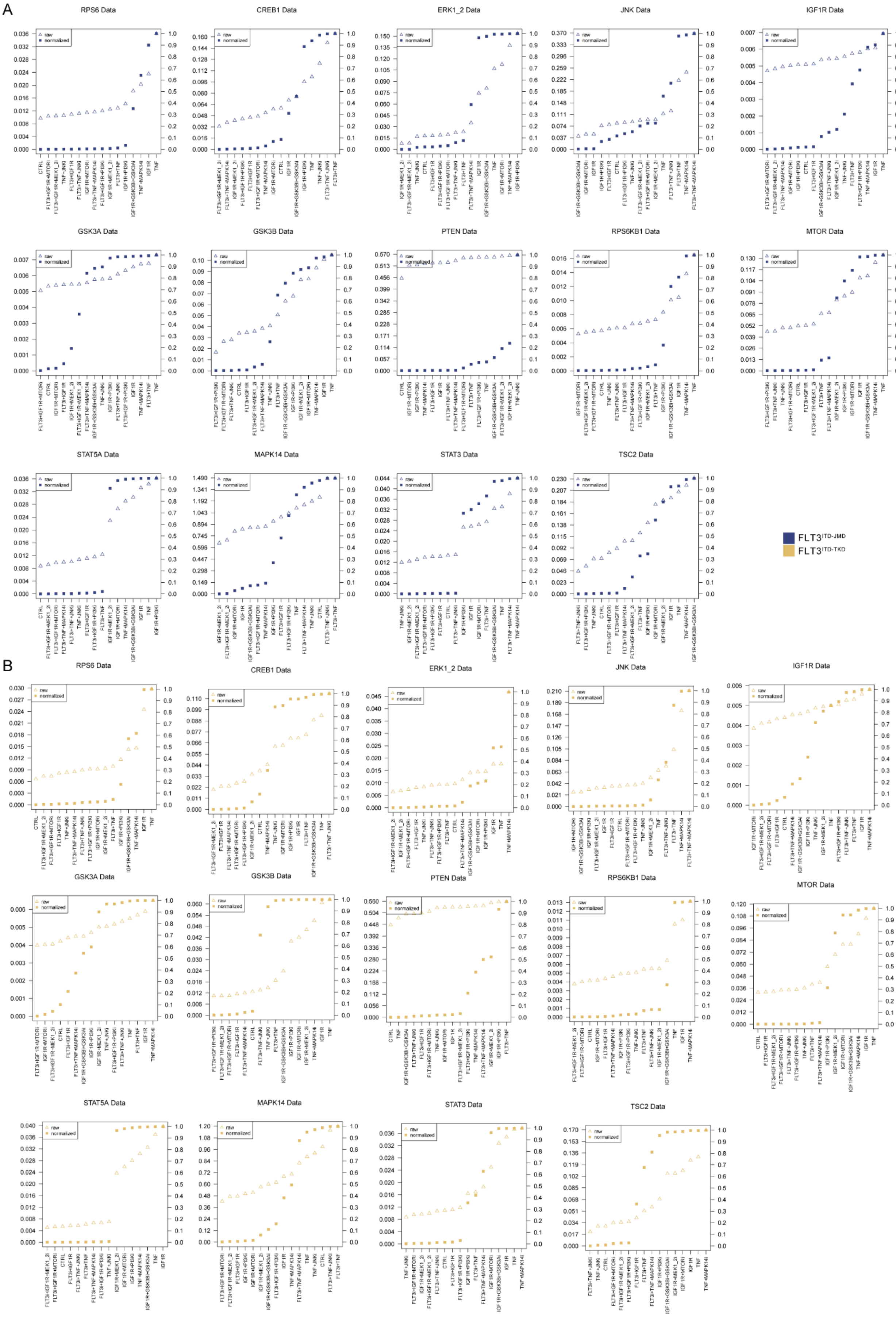
A
RPS6 Data
CREB1 Data
ERK1_2 Data
JNK Data
IGF1R Data
GSK3A Data
GSK3B Data
PTEN Data
RPS6KB1 Data
MTOR Data
STAT5A Data
MAPK14 Data
STAT3 Data
TSC2 Data
raw
normalized
FLT3ITD-JMD
FLT3ITD-TKD
B
RPS6 Data
CREB1 Data
ERK1_2 Data
JNK Data
IGF1R Data
GSK3A Data
GSK3B Data
PTEN Data
RPS6KB1 Data
MTOR Data
STAT5A Data
MAPK14 Data
STAT3 Data
TSC2 Data

**Figure S3. Normalization of analytes activity through Hill curves.** Experimental data were normalized and scaled from 0 to 1 using analyte-specific Hill functions. Raw data are reported as triangles, normalized data and squares. FLT3 $^{ITD-JMD}$ and $^{-TKD}$ plots are blue and yellow, respectively.

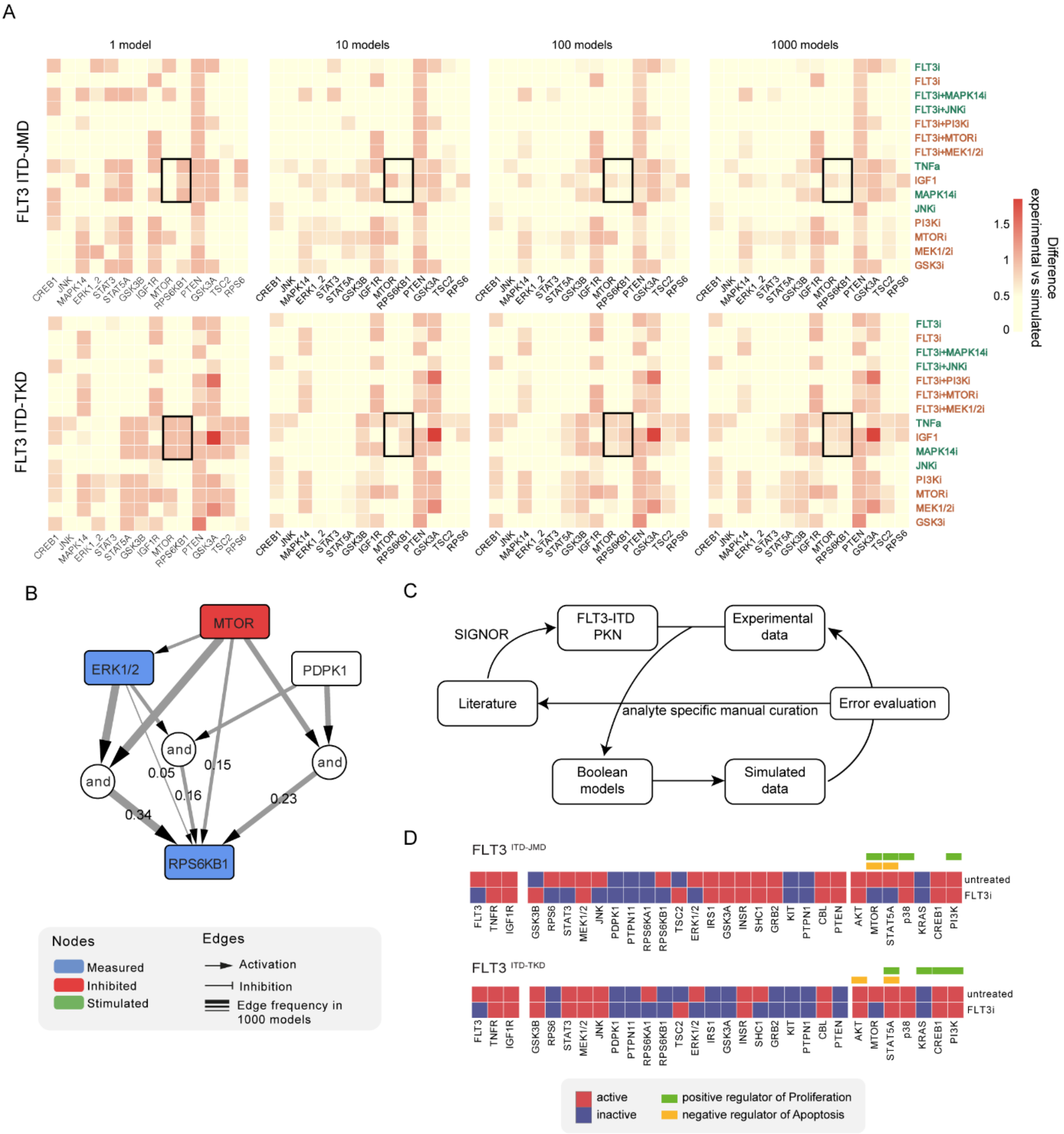
A
1 model
10 models
100 models
1000 models
FLT3 ITD-JMD
FLT3 ITD-TKD
CREB1
JNK
MAPK14
ERK1_2
STAT3
STAT5A
GSK3B
IGF1R
MTOR
RPS6KB1
PTEN
GSK3A
TSC2
RPS6
FLT3i
FLT3i
FLT3i+MAPK14i
FLT3i+JNKi
FLT3i+PI3Ki
FLT3i+MTORi
FLT3i+MEK1/2i
TNFa
IGF1
MAPK14i
JNKi
PI3Ki
MTORi
MEK1/2i
GSK3i
Difference
experimental vs simulated
1.5
1
0.5
0
B
MTOR
ERK1/2
PDPK1
and
and
and
0.05
0.15
0.16
0.34
0.23
RPS6KB1
Nodes
Measured
Inhibited
Stimulated
Edges
Activation
Inhibition
Edge frequency in
1000 models
C
SIGNOR
FLT3-ITD
PKN
Experimental
data
Literature
analyte specific manual curation
Error evaluation
Boolean
models
Simulated
data
D
FLT3 ITD-JMD
FLT3 ITD-TKD
untreated
FLT3i
FLT3
TNFR
IGF1R
GSK3B
RPS6
STAT3
MEK1/2
JNK
PDPK1
PTPN11
RPS6KA1
RPS6KB1
TSC2
ERK1/2
IRS1
GSK3A
INSR
SHC1
GRB2
KIT
PTPN1
CBL
PTEN
AKT
MTOR
STAT5A
p38
KRAS
CREB1
PI3K
active
inactive
positive regulator of Proliferation
negative regulator of Apoptosis

**Figure S4. Overview of the optimized Boolean models.**

**A**) Heatmaps represent the absolute value of the difference between model simulation results to normalized experimental data in a color-coded from light yellow to red. Simulation results of the average of respectively 1, 10, 100, and 1000 models are shown. Squared areas represent subpictures showing that a plateau in the difference is reached averaging 100 models.

**B**) The subnetwork of the regulation of RPS6KB1, derived from the FLT3 $^{ITD-TKD}$ Boolean model, shows that the family of 1000 models may contain different paths to the same node with similar frequencies.

**C**) PKN refinement cycle. We first constructed the FLT3 ITD Prior Knowledge Network (PKN) extracting the interactions from SIGNOR. Hence, the PKN was trained against the experimental data to produce the Boolean models. The simulated protein values were compared to the experimental data to evaluate the error. When we noticed that the error was systematic (e.g., the error was high for all the measurements of a specific analyte), we manually curated the interactions to improve the predictive abilities of the model.

**D)** Steady states of FLT3 $^{ITD-JMD}$ (*upper panel*) and $^{TKD}$ (*lower panel*) Boolean models in FLT3 inhibition and control conditions. Proteins are red (active) or blue (inactive). The regulators of Proliferation and Apoptosis are reported in green and orange, respectively.

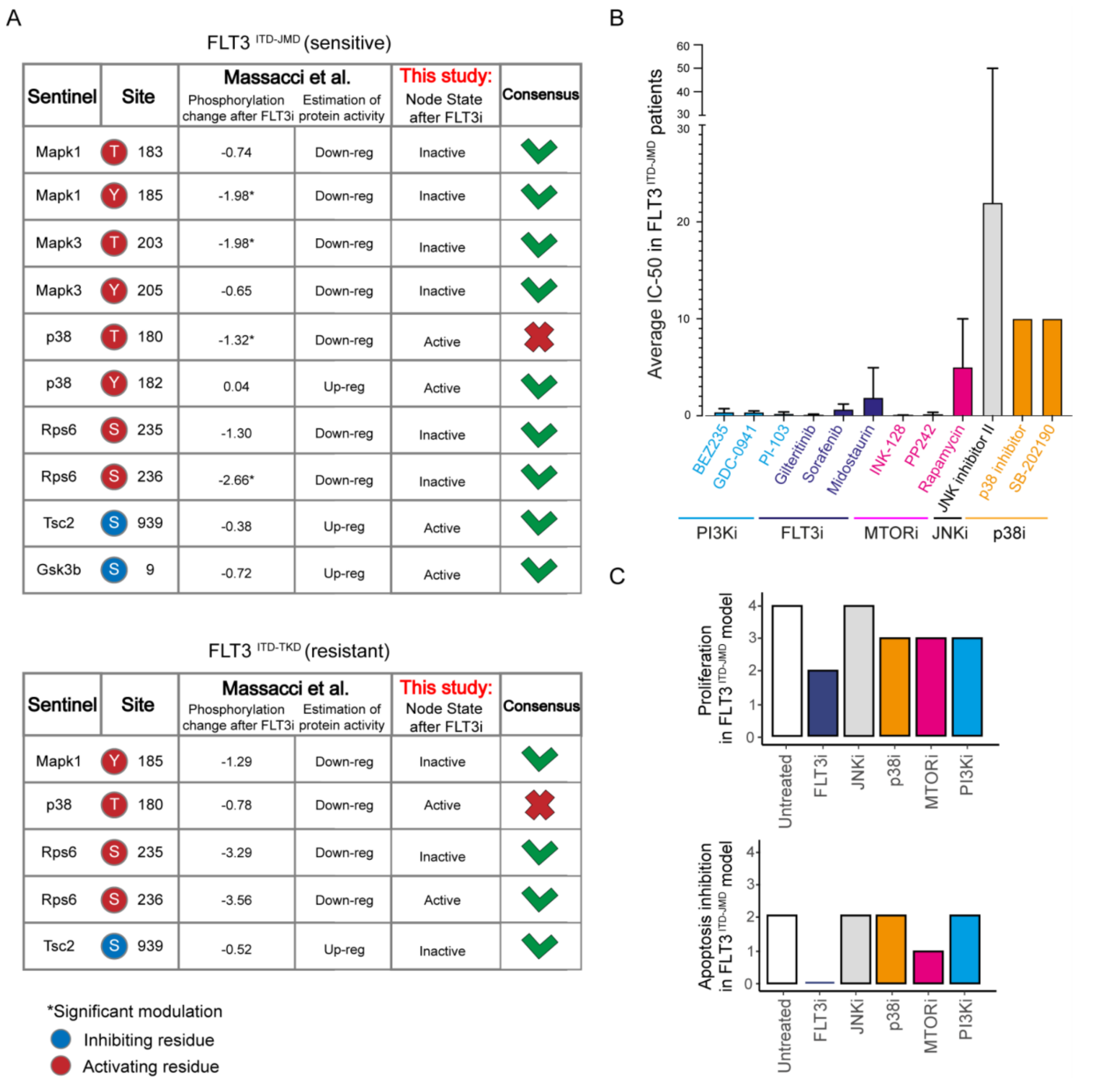

FLT3 ITD-JMD (sensitive)

| Sentinel | Site | | Massacci et al. Phosphorylation change after FLT3i | Massacci et al. Estimation of protein activity | This study: Node State after FLT3i | Consensus |
|---|---|---|---|---|---|---|
| Mapk1 | T | 183 | -0.74 | Down-reg | Inactive | ✓ |
| Mapk1 | Y | 185 | -1.98* | Down-reg | Inactive | ✓ |
| Mapk3 | T | 203 | -1.98* | Down-reg | Inactive | ✓ |
| Mapk3 | Y | 205 | -0.65 | Down-reg | Inactive | ✓ |
| p38 | T | 180 | -1.32* | Down-reg | Active | ✗ |
| p38 | Y | 182 | 0.04 | Up-reg | Active | ✓ |
| Rps6 | S | 235 | -1.30 | Down-reg | Inactive | ✓ |
| Rps6 | S | 236 | -2.66* | Down-reg | Inactive | ✓ |
| Tsc2 | S | 939 | -0.38 | Up-reg | Active | ✓ |
| Gsk3b | S | 9 | -0.72 | Up-reg | Active | ✓ |

FLT3 ITD-TKD (resistant)

| Sentinel | Site | | Massacci et al. Phosphorylation change after FLT3i | Massacci et al. Estimation of protein activity | This study: Node State after FLT3i | Consensus |
|---|---|---|---|---|---|---|
| Mapk1 | Y | 185 | -1.29 | Down-reg | Inactive | ✓ |
| p38 | T | 180 | -0.78 | Down-reg | Active | ✗ |
| Rps6 | S | 235 | -3.29 | Down-reg | Inactive | ✓ |
| Rps6 | S | 236 | -3.56 | Down-reg | Active | ✓ |
| Tsc2 | S | 939 | -0.52 | Up-reg | Inactive | ✓ |



**Figure S5. Validation of FLT3-ITD specific models on real-world independent datasets.** **A)** Comparison of our models' steady states upon FLT3 inhibition with data from Massacci et al. paper (Massacci et al., 2023). The agreement between the activation status of nodes in the FLT3$^{ITD-JMD}$ (upper table) and FLT3$^{ITD-TKD}$ (lower table) models with level of regulatory phosphorylation in the reference dataset is shown in the Consensus column.

**B)** Bar plot reporting the average IC50 values of 134 FLT3$^{ITD-JMD}$ positive patient-derived primary blasts exposed to the indicated drug treatments **C)** Bar plots depicting the (Tyner et al., 2018; Zhang et al., 2019) *in-silico* proliferation activation (up) and apoptosis inhibition (down) levels of FLT3$^{ITD-JMD}$ model in *untreated* condition (white) and following *in silico* treatment with single kinase inhibitors tested in BEAT-AML dataset (FLT3i, JNKi, p38, MTORi and PI3Ki in blue, grey, orange, purple, sky blue, respectively).

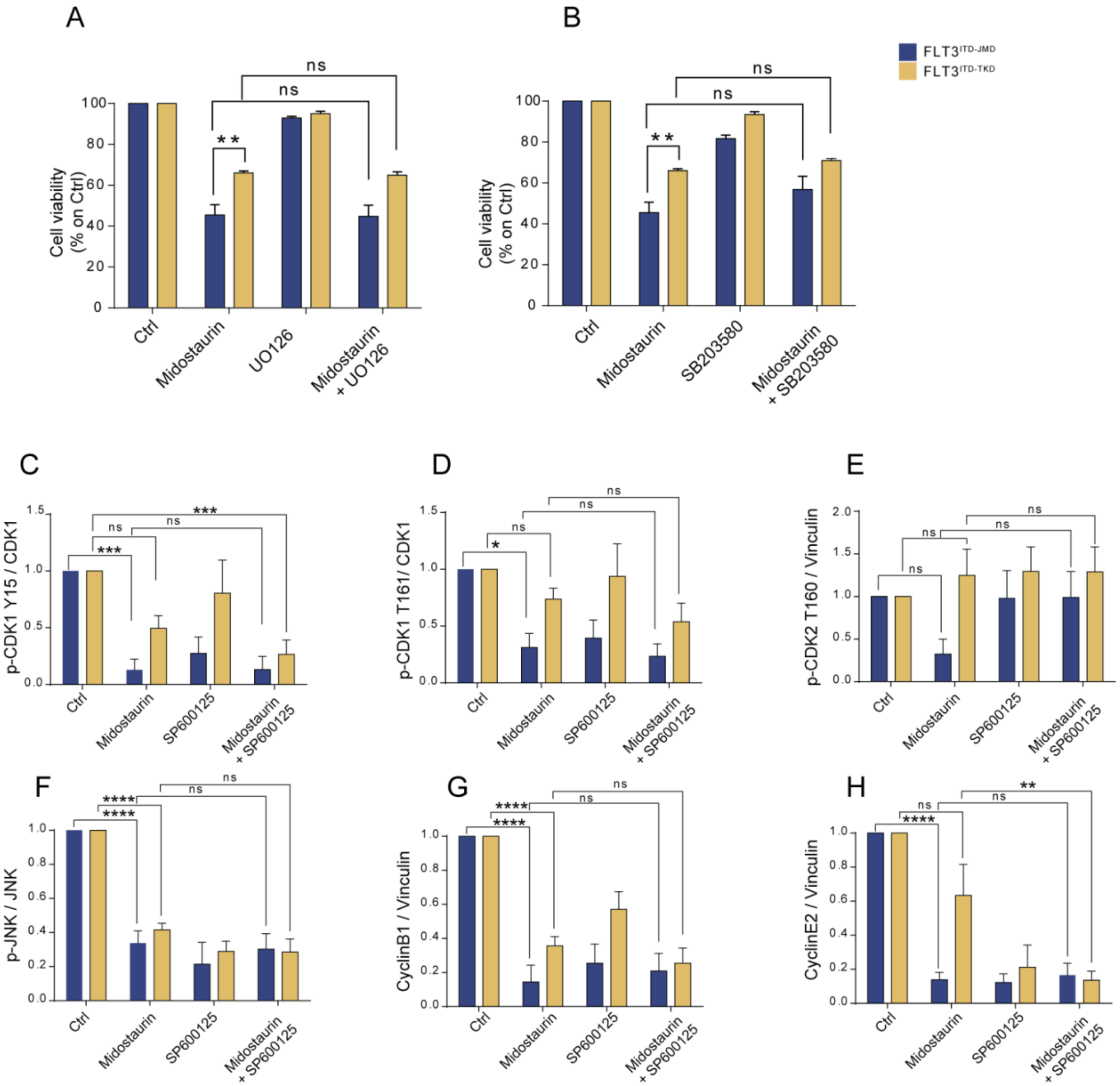


**Figure S6. Quantification of western blot analysis of cell cycle proteins.**

**A-B**) Bar plot showing the absorbance values at 595nm normalized on control conditions for FLT3 $^{ITD-JMD}$ (yellow) and $^{-TKD}$ (blue) cells treated with Midostaurin and/or UO126 (MEK inhibitor) (**A**) and with Midostaurin and/or SB203580 (p38 inhibitor) (**B**).

**C-H)** Barplots showing the densitometric quantification of the representative western blots for the proteins: phpspho-CDK1 Y15 (**C**), phospho-CDK1 T161 (**D**), phospho-CDK2 T160 (**E**), phospho-JNK (**F**), Cyclin B1 (**G**), CyclinE2 (**H**).

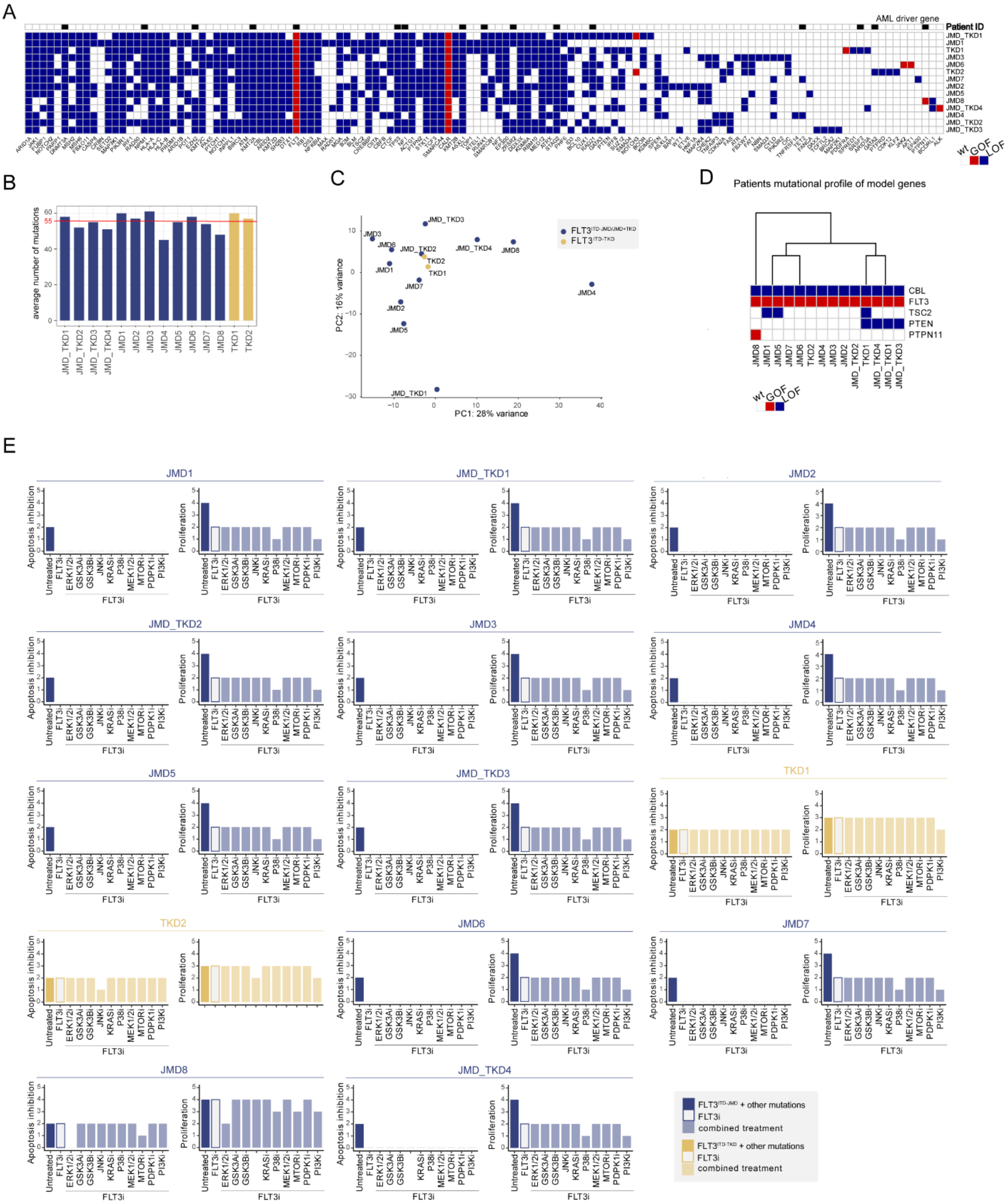
A
AML driver gene
Patient ID
wt GOF LOF
B
average number of mutations
C
PC1: 28% variance
PC2: 16% variance
D
Patients mutational profile of model genes
CBL
FLT3
TSC2
PTEN
PTPN11
E
Apoptosis inhibition
Proliferation
FLT3i
combined treatment

**Figure S7. Overview of patient-specific Boolean models.**

**A**) Heatmap showing the mutational profile (columns) of each patient (rows). AML driver genes are highlighted with black squares. White, blue and red squares represent wild type genes, 'Loss Of Function' (LOF) and 'Gain Of Function' (GOF) mutations, respectively.

**B)** Bar plot reporting the number of mutations for each patient. The red line represents the average number of mutations (55).

**C)** Principal Component Analysis (PCA) of patients' expression profile.

**D)** Heatmap reporting the mutational profile of patients restricted to genes present in the cell-derived Boolean model. White, blue and red squares represent wild type genes, 'Loss Of Function' (LOF) and 'Gain Of Function' (GOF) mutations, respectively.

**E)** Each couple of bar plots show the *in-silico* apoptosis inhibition (*right*) and proliferation activation (*left*) levels in control and FLT3 inhibition conditions in combination with the knock-out of each of 10 kinases in FLT3 $^{ITD\text{-}JMD}$ and FLT3 $^{ITD\text{-}JMD + TKD}$ (blue) and FLT3 $^{ITD\text{-}TKD}$ (blue) patients.

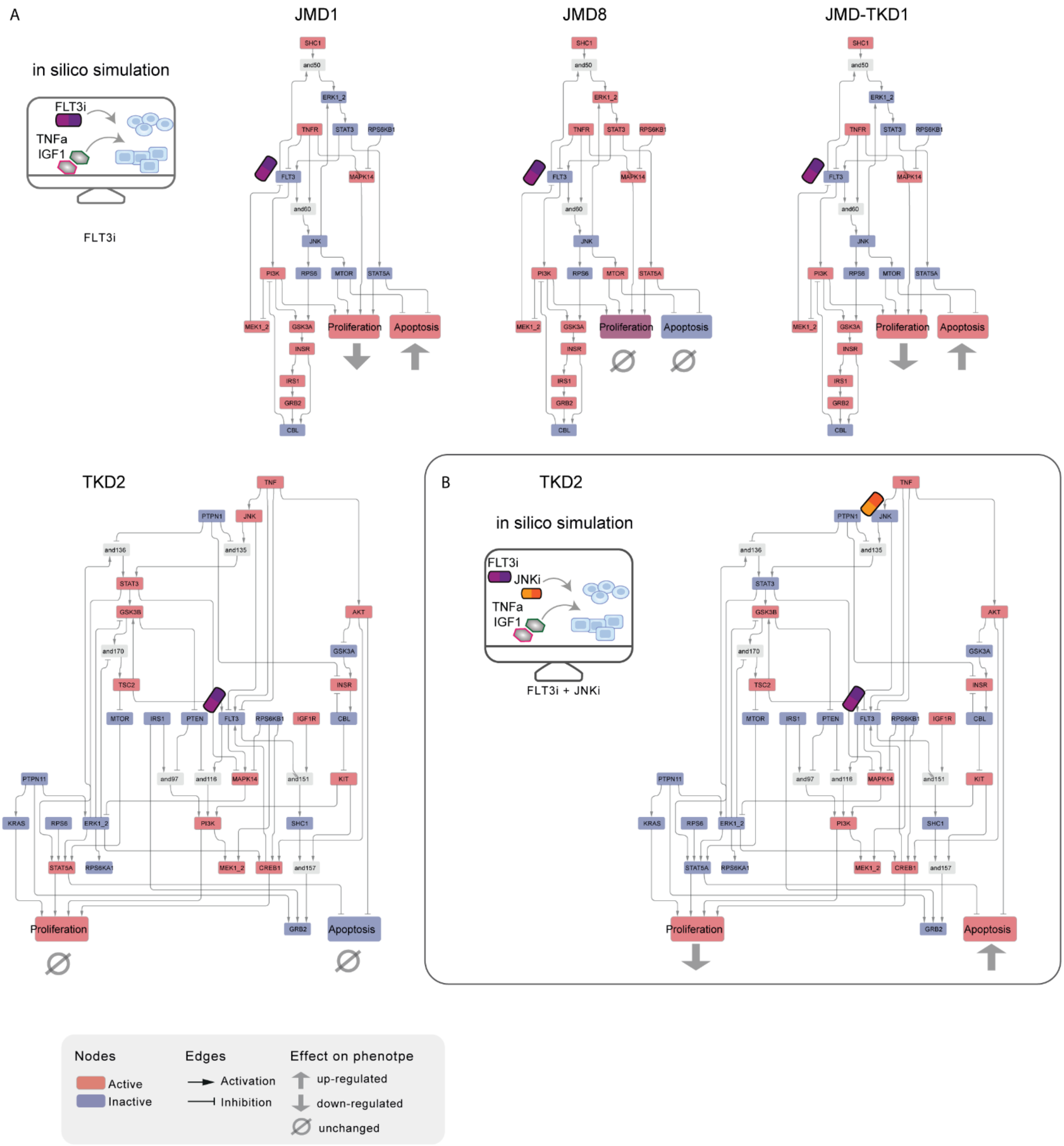


**Figure S8. Personalized Boolean models**

**A)** High-confidence Boolean models of representative patients (JMD1, JMD8, JMD-TKD1, TKD2). Nodes are color coded according to their activity upon in silico simulation of FLT3 inhibition.

**B)** High-confidence Boolean model of TKD2 patient shows how the combined inhibition of FLT3 and JNK in silico reverts the TKI resistance.

#### Supplementary Tables summary

**Table S1**: FLT3-ITD Prior Knowledge Network (PKN), table downloaded form SIGNOR, data driven edges integration, PKN in .sif format; regulators of phenotypes annotated by ProxPath resource.

**Table S2**: experimental design of the multiparametric experiment of FLT3 $^{ITD-JMD}$ and $^{ITD-TKD}$ BaF3 cell lines; treatments and analytes measured; activity readout annotation.

**Table S3**: cue-sentinel response multiparametric dataset raw data and statistics, (MILLIPLEX kit: 9plex_Cat.No.48-680MAG).

**Table S4**: cue-sentinel response multiparametric dataset, raw data and statistics (MILLIPLEX kit: 11plex_Cat.No.48-611MAG).

**Table S5**: complete cue-sentinel response multiparametric dataset used for Boolean models building in MIDAS format, raw and normalized data.

**Table S6**: data used for the validation of FLT3 ITDs Boolean models using independent resources.

**Table S7**: Panel of 262 mutations relevant to hematological malignancies analyzed in de novo AML cohort of 14 patients.

**Table S8**: results of the getITD output for the classification of the de novo AML cohort of 14 patients

**Table S9**: clinical data, mutation profile and RNAseq results of the AML patients' cohort.

#### Supplementary Materials and Methods

Here, we present a comprehensive and detailed description of the steps of our modeling strategy. Its primary objective is to provide readers with a comprehensive understanding of the rationale behind the choices made at each stage of the analysis.

The overall bioinformatics strategy was divided into two main parts:

A) Cells-specific models' generation.

B) Optimized Boolean models' validation.

C) Optimized Boolean models' usage.

For each step, we provide references to the corresponding code available in our GitHub repository (https://github.com/SaccoPerfettoLab/FLT3-ITD_driven_AML_Boolean_models). This meticulous exposition serves to enhance transparency and facilitate a more in-depth and independent assessment of our analytical procedures.

**A. Cell-specific models' generation**

To generate FLT3-ITD-specific predictive Boolean models, we exploited the genetic algorithm of CellNOptR, which trains a manually curated FLT3 Prior Knowledge Network (76 nodes and 193 edges, **Figure S1**) on perturbation data (**Table S1**).

In the first step, CellNOptR preprocesses the PKN (**Fig. S1**) and translates it into logical functions (scaffold model).

- ***Input preparation:***
    - Conversion of PKN in SIF format (**Table S1, Figure S2**).
    - Perturbation data were normalized between 0 and 1 using a customized Hill function (**Table S5, Figure S3**).

  The code to reproduce this step is available on GitHub in the A. Model generation section, Paragraph 1.

- ***Pre-processing of the Prior Knowledge Network:*** this phase is needed to achieve, from the PKN, a simplified yet informative network (here dubbed 'scaffold model'). In this phase, we remove unnecessary nodes such as chains of signaling interaction with the same sign, and we check that the scaffold model has the potential to display an output (modulation of a sentinel) for any input provided (cue). This phase accounts for three main steps:

- ***Compression:*** unmeasured and untargeted proteins are removed
- ***Expansion:*** the remaining nodes are connected to every possible combination of upstream regulators with both ‘AND’ and ‘OR’ Boolean operators. The ‘AND’ operators are added as nodes of the network. Thus, we end with 204 nodes (30 proteins and 174 AND Boolean operators) and 612 edges.
- ***Imputation:*** This step is crucial to add co-regulations that may happen in our system but are still not known or annotated in literature and thus are missing in the PKN. The R package *CNORfeeder* was exploited to integrate the interactions derived from correlation among analytes in the perturbation data. This step allowed us to add 144 data-derived edges in the $FLT3^{ITD\text{-}JMD}$ model and 170 in the $FLT3^{ITD\text{-}TKD}$.

Using this strategy, we obtained two FLT3-ITD specific scaffold models, accounting for 206 and 208 nodes and 756 and 782 edges, for $FLT3^{ITD\text{-}JMD}$ and $FLT3^{ITD\text{-}TKD}$ respectively. The code to reproduce this step is available in our GitHub repository in the A. Model generation section, Paragraph 2.

- ***Network model optimization:*** in this step, we took advantage of a successful strategy used in our previous work (Sacco et al., 2012). The aim of this step is to train and optimize a Boolean logic model able to reproduce in silico our cue–sentinel–response multiparametric training dataset (**Fig. 3A**). In this phase, we followed standard practice in logical modeling (Dorier et al., 2016; Traynard et al., 2017):

  - *Generation of the family of 1000 Boolean models:* we run the genetic algorithm of CellNOptR package 1000 times to generate a family of 1000 optimized Boolean models for each cell line. Briefly, this choice is driven by:
    - the stochastic nature of the genetic algorithm.
    - the possibility of obtaining quantitative predictions averaging the discrete node state in each model. This feature offers the opportunity to readily evaluating the optimization process, since the agreement between experimental and simulated data increases when the predictions are carried out by averaging a larger number of models.

- *Selection of 100 optimal models:* given the observation that the agreement between experimental and simulated data reached an apparent plateau at 100 models (**Figure S4**), we filtered out only the 100 models that better fit the experimental data (Dorier et al., 2016). Then, for each cell line, we compared the experimental activity modulation of each protein (**Figure 3A, panel 1**) with the average activity modulation of the protein in the family of 100 models (**Figure 3A, panel 2**). The fit between simulated and experimental data is reported in **Figure 3A, panel 3**.

The code to reproduce this step is available in our GitHub repository in the A. Model generation section, Paragraph 3.

Importantly, the performance of the model strongly depends on the topology of the PKN. In compliance with standard practices in logical modelling (Dorier et al., 2016; Traynard et al., 2017) we performed several rounds of PKN check and adjustment, and, in each round, the entire process was iterated until the simulation provided the best fit of the available data (**Fig. S4B-C**).

- ***Visualization of optimized models:***
  - *Final model selection:* to perform further analyses we selected only the best model. The strength of this approach is that Boolean rules are derived from the combination of prior knowledge and experimental data. However, the optimization process creates a family of Boolean models where some nodes are regulated by different and opposite Boolean rules making it difficult to perform simulations (**Figure S4B**). To resolve this ambiguity and have a single Boolean rule for each node and, hence, to exploit the intrinsic predictive power of Boolean models, we selected the model with the lowest error between experimental and simulated data in $FLT3^{ITD\text{-}JMD}$ and $FLT3^{ITD\text{-}TKD}$ cell lines (best model).

    The so-obtained final models consist of 68 and 60 nodes (of which 38 and 30 are AND operators) and 161 and 133 edges for $FLT3^{ITD\text{-}JMD}$ and $FLT3^{ITD\text{-}TKD}$, respectively.
  - *High confidence model generation:* to keep a measure of reliability from the whole optimization procedure in each best model, we added the frequency of each edge in the 100 models as an attribute in each cell-specific best model. The

two final Boolean models with the highest edge confidence (frequency in 100 models > 0.4) are shown in **Figures 3B and 3C**.

The code to reproduce these steps is available in our GitHub repository in the B. Model visualization.

**B. Assessment of optimized models**

At this point, we aimed to assess whether the newly generated FLT3$^{ITD-JMD}$ and FLT3$^{ITD-TKD}$ Boolean models could recapitulate *in silico* the TKI-induced modulation of apoptosis and proliferation. To this aim, we defined a strategy that is composed of three steps:

1. We set up two initial conditions of the models aiming at reproducing the biological context of our cells before and after the FLT3 inhibitor treatment.
2. We computed the steady state of each Boolean model to derive the states of the model's proteins associated to each initial condition.
3. We unbiasedly associated the final conditions to phenotypic states of the cells.

We here in brief describe the rationale behind these choices and relative technical details.

*1. Initial conditions of the simulation*

Briefly, we defined *"Untreated"* the condition that reproduce a malignant state where cells proliferate and escape apoptosis. From the model perspective this condition is represented by an active state of the FLT3, IGF1R, and TNFR nodes.

Next, we defined *"FLT3i"* the condition that reproduces malignant cells treated with the FLT3 inhibitor (Midostaurin). From the model perspective this condition is represented by an active state of the IGF1R, and TNFR nodes and an inactive state of FLT3.

Summing up, the conditions used were:

- *Untreated condition,* in which all the receptors included in the model (FLT3, IGF1R, and TNFR) are set to ON.
- *FLT3 inhibition condition (FLT3i),* in which FLT3 was set to OFF.

*2. Computation of steady states:* here we used the *simulatorT1* function of CellOptR package, which performs a Synchronous Boolean simulation to compute the steady state of each cellular model in two conditions (**Figure S4D**).

*3. Apoptosis and proliferation activity inference:* to functionally interpret the results of the simulations, we derived the levels of 'apoptosis inhibition' and 'proliferation activation' in each condition (**Figure 3E**). To do that:

- We annotated all the proteins in the network as activators or inhibitors of 'apoptosis' and 'proliferation' phenotypes using our recently published research ProxPath (Iannuccelli et al., 2022).
- For the inference of phenotypes, we considered only proteins that were (i) regulators of a phenotype and (ii) endpoint proteins in high-confidence signaling axes (edge frequency 0.4, **Figure 3B e 3C**). This choice is driven by the need to avoid redundancy and consider the final nodes of the model which are the real effector proteins according to our mutations-specific FLT3 Boolean models.
- Then, we integrated the signal of the phenotype regulators proteins (**Figure 3E** heatmaps) to compute the level of 'apoptosis inhibition' and 'proliferation activation' in each cell line (**Figure 3E** barplot). Integration, in this context, means computing the sum of the 'scores' of proteins that influence each phenotype, with an underlying assumption of equal importance for both inhibitors and activators (OR logic).

As a result of this step, we obtained a comprehensive model encompassing proteins and phenotypes (apoptosis and proliferation) that can possess multiple values, effectively creating a multi-valued model. This enhancement enables a finer comparison between different cell lines.

The code that describes this step is available in our GitHub repository in the C. In silico validation section.

**C. Optimized boolean models' usage**

Given the ability of the ability of FLT3$^{ITD-JMD}$ and FLT3$^{ITD-TKD}$ networks to reproduce the different sensitivity to TKIs, the generated ITD-specific Boolean models were exploited to find a co-treatment, in addition to FLT3 inhibition, that could revert the resistant phenotype.

For each cell line, we inferred the levels of 'apoptosis inhibition' and 'proliferation activation' upon every possible combinate inhibition of FLT3 and key signaling kinases (ERK1/2, MEK1/2, GSK3A/B, IGF1R, JNK, KRAS, MEK1/2, MTOR, PDPK1, PI3K) in the model (**Figure 4A**). Briefly, here we used the same strategy described in "***Assessment of optimized models"*** section. The initial conditions considered were:

- *Untreated,* in which all the receptors included in the model (FLT3, IGF1R, and TNFR) are set to ON.
- *FLT3i,* in which FLT3 was set to OFF.
- *Combinatory treatment* (e.g., FLT3i + KRASi), in which both FLT3 and the target of the specific inhibition (in this case KRAS) were set to OFF.

The code to reproduce this step is available in our GitHub repository in the D. Combinatory treatment inference.
